# Misaligned chromosomes that satisfy the spindle assembly checkpoint are a strong predictor of micronuclei formation in dividing cancer cells

**DOI:** 10.1101/2022.07.20.500828

**Authors:** Ana Margarida Gomes, Bernardo Orr, Marco Novais-Cruz, Filipe De Sousa, Cristina Ferrás, Helder Maiato

**Affiliations:** Chromosome Instability & Dynamics Group, i3S - Instituto de Investigação e Inovação em Saúde, Universidade do Porto, Rua Alfredo Allen 208, 4200-135 Porto, Portugal; Instituto de Biologia Molecular e Celular, Universidade do Porto, Rua Alfredo Allen 208, 4200-135 Porto, Portugal; Cell Division Group, Department of Biomedicine, Faculdade de Medicina, Universidade do Porto, Alameda Prof. Hernâni Monteiro, 4200-319 Porto, Portugal

**Keywords:** mitosis, micronuclei, kinetochore, Mad2, Cyclin B1, chromosome congression, spindle assembly checkpoint, aneuploidy, chromosomal instability, cancer

## Abstract

Chromosome alignment to the spindle equator is a hallmark of mitosis that is thought to promote chromosome segregation fidelity in metazoans. Yet, chromosome alignment is only indirectly supervised by the spindle assembly checkpoint (SAC) as a byproduct of chromosome bi-orientation, and the consequences of defective chromosome alignment remain unclear. Here we investigated how human cells respond to chromosome alignment defects of distinct molecular nature by following the fate of live HeLa cells after RNAi-mediated depletion of 120 proteins previously implicated in chromosome alignment. Surprisingly, in all cases, cells frequently entered anaphase after a delay with chronically misaligned chromosomes. Using depletion of key proteins as prototypes for defective chromosome alignment, we show that chronically misaligned chromosomes often satisfy the SAC and directly missegregate. In-depth analysis of specific molecular perturbations that prevent proper kinetochore-microtubule attachments revealed that chronically misaligned chromosomes that missegregate frequently result in micronuclei. Higher-resolution live-cell imaging indicated that, contrary to most anaphase lagging chromosomes that correct and reintegrate the main nuclei, chronically misaligned chromosomes are a strong predictor of micronuclei formation in a cancer cell model of chromosomal instability, but not in normal near-diploid cells. We provide evidence supporting that intrinsic differences in kinetochore-microtubule attachment stability on misaligned chromosomes account for this distinct outcome. Thus, chronically misaligned chromosomes that satisfy the SAC may represent a previously overlooked mechanism driving chromosomal/genomic instability during cancer cell division, and we unveil genetic conditions predisposing for these events.

## Introduction

Chromosome alignment to the spindle equator (also known as chromosome congression) is a hallmark of mitosis that promotes chromosome segregation fidelity in metazoans by preventing chromosome dispersion upon sister chromatid separation during anaphase ^1–3^. Depending on the initial position of each chromosome at nuclear envelope breakdown (NEB), chromosome alignment in human cells can be achieved by at least two mechanisms (reviewed in ^4^). Those chromosomes that are favorably located between two newly formed spindle poles have a greater chance to become bi-oriented, with kinetochores on each chromatid attaching to microtubules oriented to opposite poles. In contrast, more peripheral chromosomes are first brought to the vicinity of spindle poles along laterally attached astral microtubules by the microtubule minus-end directed motor protein dynein at kinetochores ^5–7^, and subsequently transported towards the equator by the microtubule plus-end directed kinesin motor CENP-E ^8, 9^. Regulation of kinetochore motors with opposite activities is controlled by microtubule detyrosination, which inhibits dynein-mediated transport, while favoring CENP-E-mediated alignment of chromosomes ^10, 11^. Thus, chromosome alignment in human cells relies on the concerted action of motor-dependent and independent mechanisms, which are determined by chromosome positioning at NEB, the establishment of end-on or lateral kinetochore-microtubule interactions and specific tubulin post-translational modifications.

Despite its key role in promoting mitotic fidelity, chromosome alignment is only indirectly supervised by the spindle assembly checkpoint (SAC), which monitors the establishment of end-on kinetochore-microtubule attachments required for chromosome bi-orientation and regulates the metaphase-anaphase transition ^12, 13^. It is therefore widely assumed that, under physiological conditions, cells only enter anaphase once all chromosomes align and bi-orient ^14–22^. However, chromosome alignment may occur independently of end-on kinetochore-microtubule attachments and chromosome bi-orientation ^8, 23, 24^, and conditions exist in which vertebrate cells enter anaphase in the presence of misaligned chromosomes. For example, approximately 10% of newt lung cells in culture enter anaphase in the presence of one or more misaligned chromosomes ^25^. Additionally, misaligned chromosomes generated after functional perturbation of the kinetochore-associated CENP-E/Kinesin-7 in primary mouse fibroblasts and human HeLa cells in culture, as well as in regenerating hepatocytes in vivo, did not prevent anaphase onset in approximately 25%, 40% and 95% of cell divisions, respectively, resulting in aneuploidy ^26–28^. Importantly, as opposed to massive aneuploidy that renders cells unviable and has a tumor suppressing effect ^29, 30^, gain/loss of just one or few chromosomes that are unable to complete alignment represents a real threat to chromosomal stability and has been shown to contribute to tumorigenesis in vivo ^31^. Thus, understanding how human cells respond to chromosome alignment defects and determining what happens to a chronically misaligned chromosome remain fundamental unanswered questions with strong clinical implications.

Here we used high-content live-cell imaging combined with RNAi of 120 proteins previously implicated in chromosome alignment to inquire how human cells respond to chromosome alignment defects of distinct molecular nature. This allowed a thorough investigation of the consequences of a broad range of chromosome alignment defects for mitotic fidelity and cell viability. Surprisingly, we found that, regardless of the underlying molecular perturbation, cells frequently entered anaphase after a delay with misaligned chromosomes. Complementary experiments focusing on a subset of genes required for proper kinetochore-microtubule attachments and chromosome bi-orientation indicate that chronically misaligned chromosomes often satisfy the SAC and missegregate, leading to the formation of micronuclei. Higher resolution live-cell microscopy analysis revealed that entry into anaphase with misaligned chromosomes is a frequent outcome and the most faithful predictor of micronuclei formation, specifically during division of unperturbed chromosomally unstable cancer cells. We provide data suggesting that intrinsic differences in kinetochore-microtubule attachment stability relative to normal near-diploid cells account for this outcome. Thus, in addition to anaphase lagging chromosomes, the present work identifies an alternative pathway that may drive chromosomal instability and account for micronuclei formation during cancer cell division.

## Results

### A broad range of chromosome alignment defects directly lead to missegregation

To systematically inquire how human cells respond to chromosome alignment defects of distinct molecular nature we used siRNAs to knockdown 120 different proteins previously implicated in this process (Table S1 and S2, Figure S1 and S2), combined with high-content live-cell microscopy in human HeLa cells stably expressing histone H2B-GFP (to visualize chromosomes) and α-tubulin-mRFP (to visualize mitotic spindles) (Figure 1). Under our imaging conditions HeLa cells treated with a control scrambled siRNA underwent consecutive rounds of mitosis and completed chromosome alignment in 23±8 min (mean ± s.d., n=7229 cells), indicating no relevant phototoxicity (Movie S1). In contrast, cells treated with siRNAs against target proteins previously implicated in chromosome alignment, consistently showed three main mitotic phenotypes: 1) cells that entered anaphase after a delay in completing chromosome alignment (≥2 s.d. in control-depleted cells); 2) cells that entered anaphase without completing chromosome alignment; and 3) cells that died in mitosis without completing chromosome alignment (Figure 2a, b). In some conditions, such as depletion of several Augmin complex subunits ^32^, CLASPs ^33^ or the Ska complex ^34^, among others, a fraction of cells was also unable to maintain chromosome alignment after completing congression to the spindle equator and showed signs that resembled cohesion fatigue and/or loss of spindle pole integrity (Figure S3a, b, and Movie S2). Not surprisingly, defective chromosome alignment was often associated with a significant mitotic delay, indicating a functional SAC whose satisfaction was nevertheless compromised (Figure 2b and Figure S4). Moreover, the severity of the observed chromosome alignment defects varied extensively, suggesting that certain proteins are more crucial for this process than others (Figure 2b). This was the case of the kinetochore-associated kinesin CENP-E and its associated protein TRAMM ^35^, several cytoplasmic Dynein subunits, members of the KNL1, Mis12 and Ndc80 (KMN) network ^36^, the Ska complex ^34^, and the Augmin complex ^32^ (Movie S2, S3 and S4). However, it remains possible that less penetrant phenotypes were due to sub-optimal protein depletion. Nevertheless, and regardless of phenotypic penetrance, it was surprising that in all the different molecular perturbations performed, cells frequently entered anaphase with chronically misaligned chromosomes that often missegregate.

**Figure 1.**
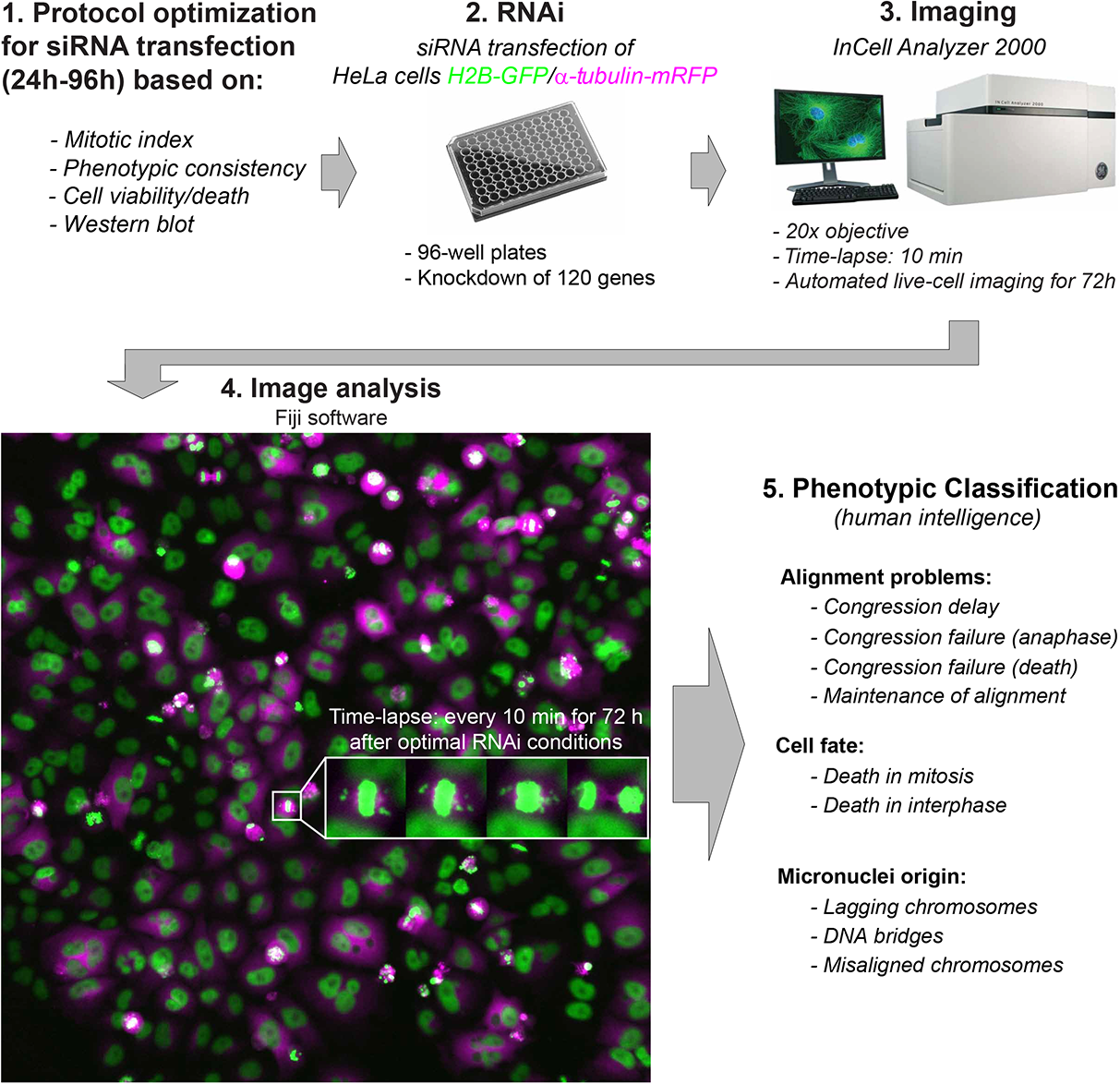
Schematic illustration of the high-content analysis of chromosome alignment defects. Different steps between protocol optimization and automated live-cell imaging of 120 different RNAi conditions against genes previously implicated in chromosome congression in HeLa cells stably expressing H2B-GFP and α-tubulin-mRFP to visualize chromosomes and microtubules, respectively. An example of a typical image gallery from the time-lapse movies is shown, highlighting one particular cell with congression failure. The distinct phenotypes analyzed are indicated.

**Figure 2.**
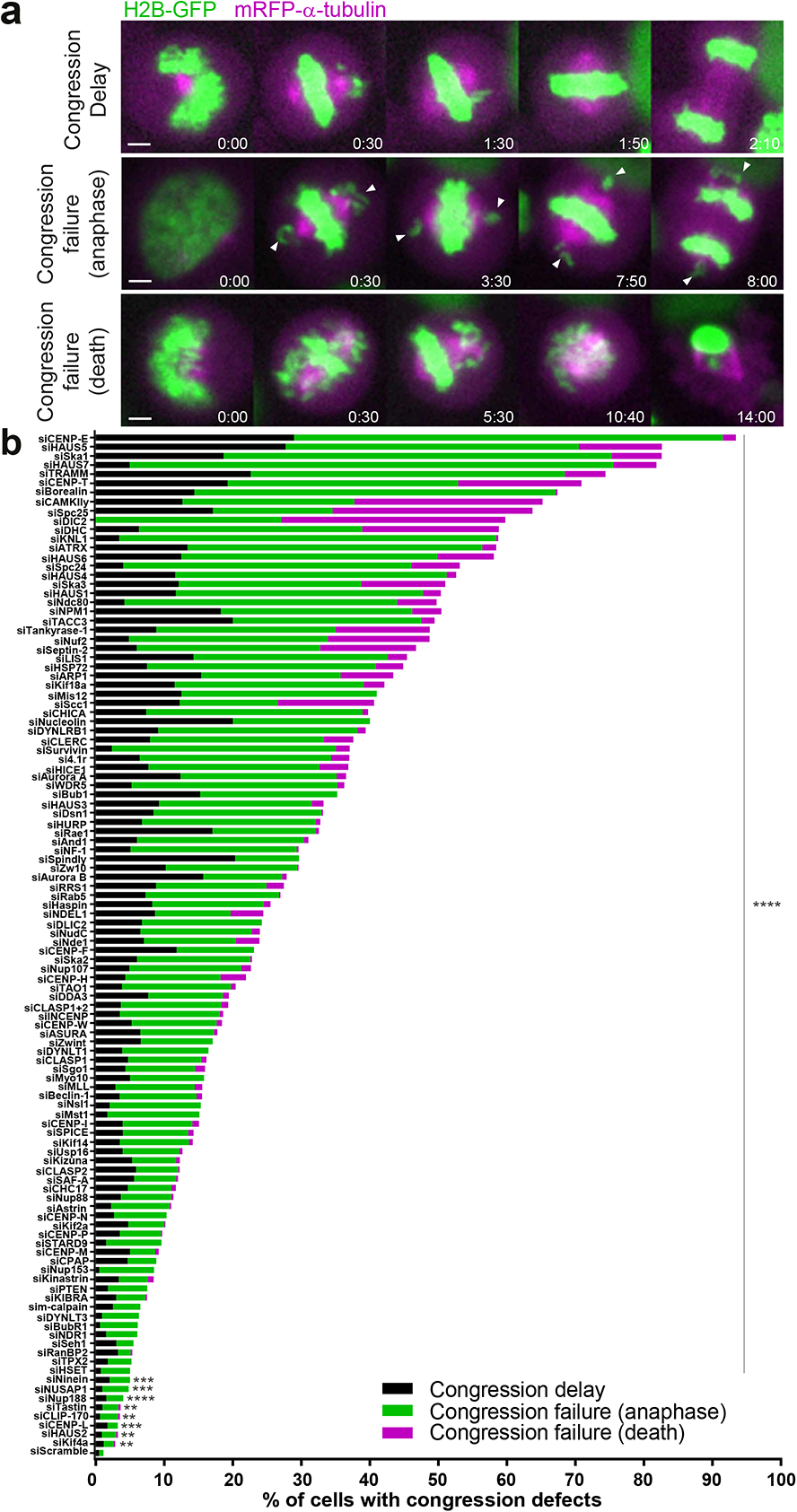
A broad range of chromosome alignment defects directly lead to missegregation. **a)** Examples of time-lapse sequences illustrating the three main mitotic phenotypes observed. Arrows indicate chromosomes at the poles in cells exhibiting chromosome alignment defects. Pixels were saturated optimal visualization of misaligned chromosomes. Scale bar: 5 µm. Time: h:min. **b)** Quantification of congression phenotypes in control (siScramble) and siRNA-depleted cells. At least 2 independent experiments per condition were performed. The total number of cells analyzed for each condition is indicated in Table S1. (*p≤0.05, **p≤0.01, ***p≤0.001, ****p≤0.0001, ns corresponds to not significantly different from control; Fisheŕs exact two-tailed test).

### Mild, yet penetrant, chromosome alignment defects are compatible with mitotic progression and cell viability

Next, we investigated how the extent of chromosome alignment defects impacts cell viability during and after mitosis (Figure 3a, b). We found a positive correlation (r >0.5) between the propensity of cells to die in mitosis and the severity of the chromosome alignment defects that typically associate with a stronger mitotic delay (Figure 3b-d). A positive, yet weaker, correlation (r >0.3) was also observed between the likelihood of cells to die in the subsequent interphase and the time they spent in mitosis due to chromosome alignment defects (Figure 3b, e, f). In one particular case (NUP107 RNAi), most cells died in the subsequent interphase likely due to a well-established role in nuclear pore complex assembly and function (reviewed by ^37^). Interestingly, some conditions that led to a high frequency of cells with congression problems, such as CENP-E or Kif18a depletion, in which cells entered anaphase with only few misaligned chromosomes or a less compact metaphase plate, respectively (Movie S5) (see also ^2, 26, 28, 38^), did not result in a proportional increase in cell death (Figure 3b). Thus, severe defects in chromosome alignment that prevent timely SAC satisfaction interfere with cell fate, ultimately compromising cell viability during mitosis and/or in the subsequent interphase. However, certain molecular perturbations that result in milder, yet highly penetrant, congression defects, represent exceptions to this rule and are compatible with mitotic progression and cell viability, thereby representing a threat to chromosomal stability.

**Figure 3.**
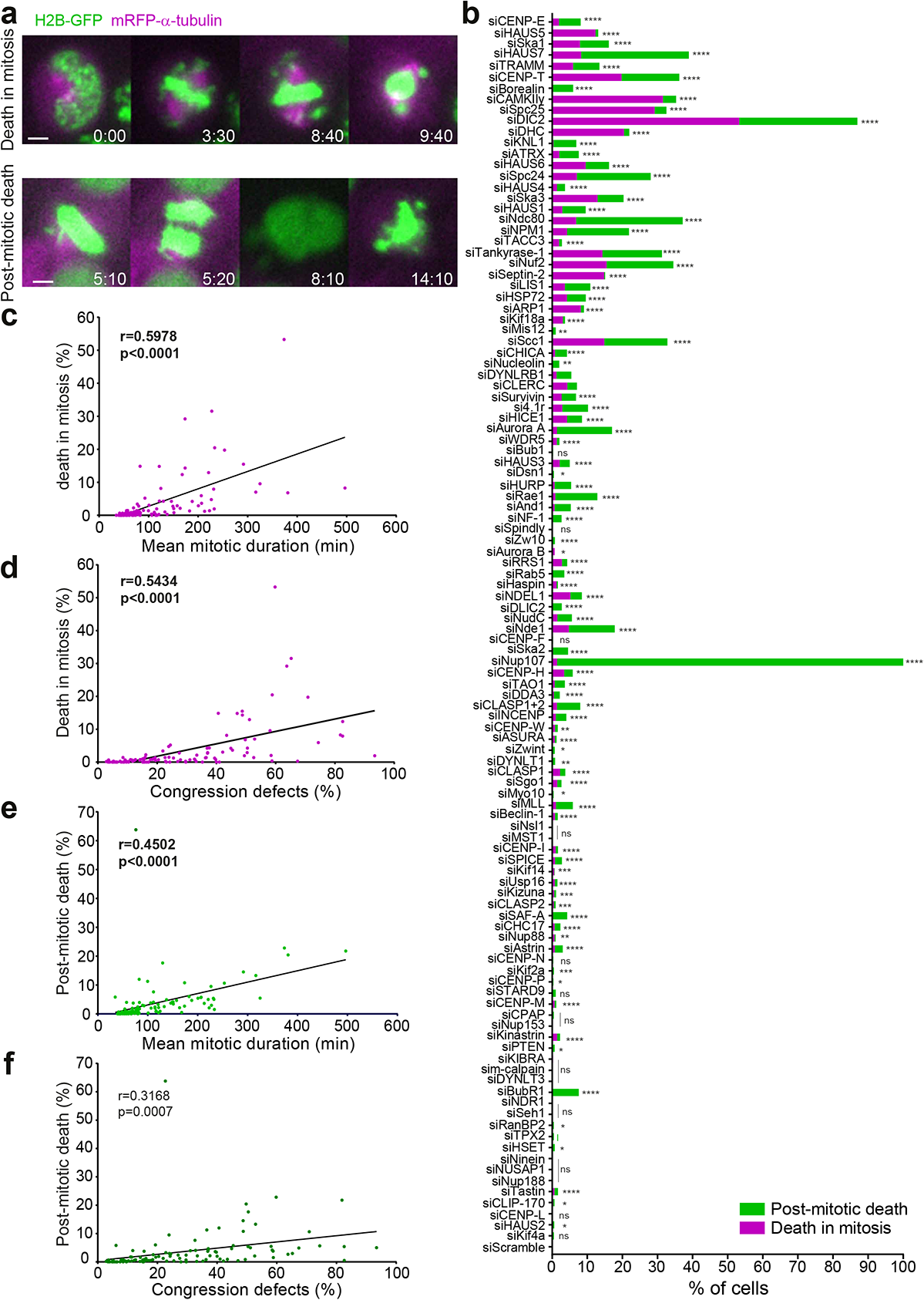
Mild, yet penetrant, chromosome alignment defects are compatible with mitotic progression and cell viability. **a)** Examples of time-lapse sequences illustrating the fates exhibited by HeLa cells undergoing congression defects following siRNA knockdown. Some cells died in mitosis, while others exited mitosis after a prolonged mitotic delay and died in interphase. Time: h:min, from nuclear envelope breakdown (NEB) to each cellular outcome. Scale bar: 5 µm. **b)** Frequency of cells that either died in mitosis (magenta) or died in interphase (green) in control and siRNA-depleted cells. The total number of cells analyzed for each condition is indicated in Table S1. (*p≤0.05, **p≤0.01, ***p≤0.001, ****p≤0.0001, ns corresponds to not significantly different from control; Fisheŕs exact two-tailed test). **c)** Correlation between the mean mitotic duration and the percentage of cell death in mitosis for each condition. **d)** Correlation analysis was performed to analyze the correlation between the severity of the phenotypes and the frequency of cell death in mitosis. **e)** Correlation between the mean mitotic duration after siRNA treatment and the percentage of cells that died in interphase. **f)** Correlation between the level of severity of the phenotypes and the frequency of cell death in interphase. Plots show data obtained from live-cell imaging after the indicated siRNA treatment conditions, analyzed using two-tailed test. Pearson’s correlation (r) and respective P-values are indicated in the plots.

### Cells with chronically misaligned chromosomes enter anaphase after satisfying the spindle assembly checkpoint and undergoing normal Cyclin B1 degradation

Given the high risk of chromosome missegregation in cells that enter anaphase with chronically misaligned chromosomes, we investigated whether they do so after SAC satisfaction or by bypassing an active SAC, a process known as mitotic slippage ^39^. In contrast to cells that satisfy the SAC, human cells undergoing mitotic slippage upon complete microtubule depolymerization with nocodazole retain the SAC proteins, Mad1, Mad2 and BubR1 at kinetochores and very slowly degrade Cyclin B1 due to residual APC/C activity ^40–42^. To distinguish between these possibilities, we used live imaging in HeLa cells stably expressing Mad2-GFP to monitor the status of the SAC in control- or CENP-E-depleted cells that enter anaphase with one or few chronically misaligned chromosomes at very high frequency (Figure 2b; see also ^27, 28^). As expected, in cells treated with a control siRNA Mad2-GFP accumulated at kinetochores during prometaphase and gradually disappeared as chromosomes bi-oriented and aligned at the metaphase plate, being undetectable at kinetochores when cells entered anaphase (Figure 4a and Movie S6). Likewise, Mad2-GFP accumulated exclusively at the kinetochores from those few chromosomes that never complete alignment after CENP-E depletion, becoming undetectable before anaphase onset and throughout anaphase (Figure 4a and Movie S6). To obtain a more quantitative picture, we used immunofluorescence in fixed HeLa cells to measure the fluorescence of the SAC protein Mad1 relative to CENP-C (a constitutive kinetochore component) on persistently misaligned chromosomes after CENP-E depletion in early anaphase (Figure 4b). We found that, in striking contrast to misaligned chromosomes during prometaphase where Mad1 signal was clearly detected at kinetochores in both control- and CENP-E-depleted cells (Figure 4c), virtually no Mad1 signal was detected at both kinetochores from misaligned chromosomes that persisted in early anaphase after CENP-E depletion (Figure 4c). Together, these data suggest that cells with chronically misaligned chromosomes enter anaphase after a delay by satisfying the spindle assembly checkpoint.

**Figure 4.**
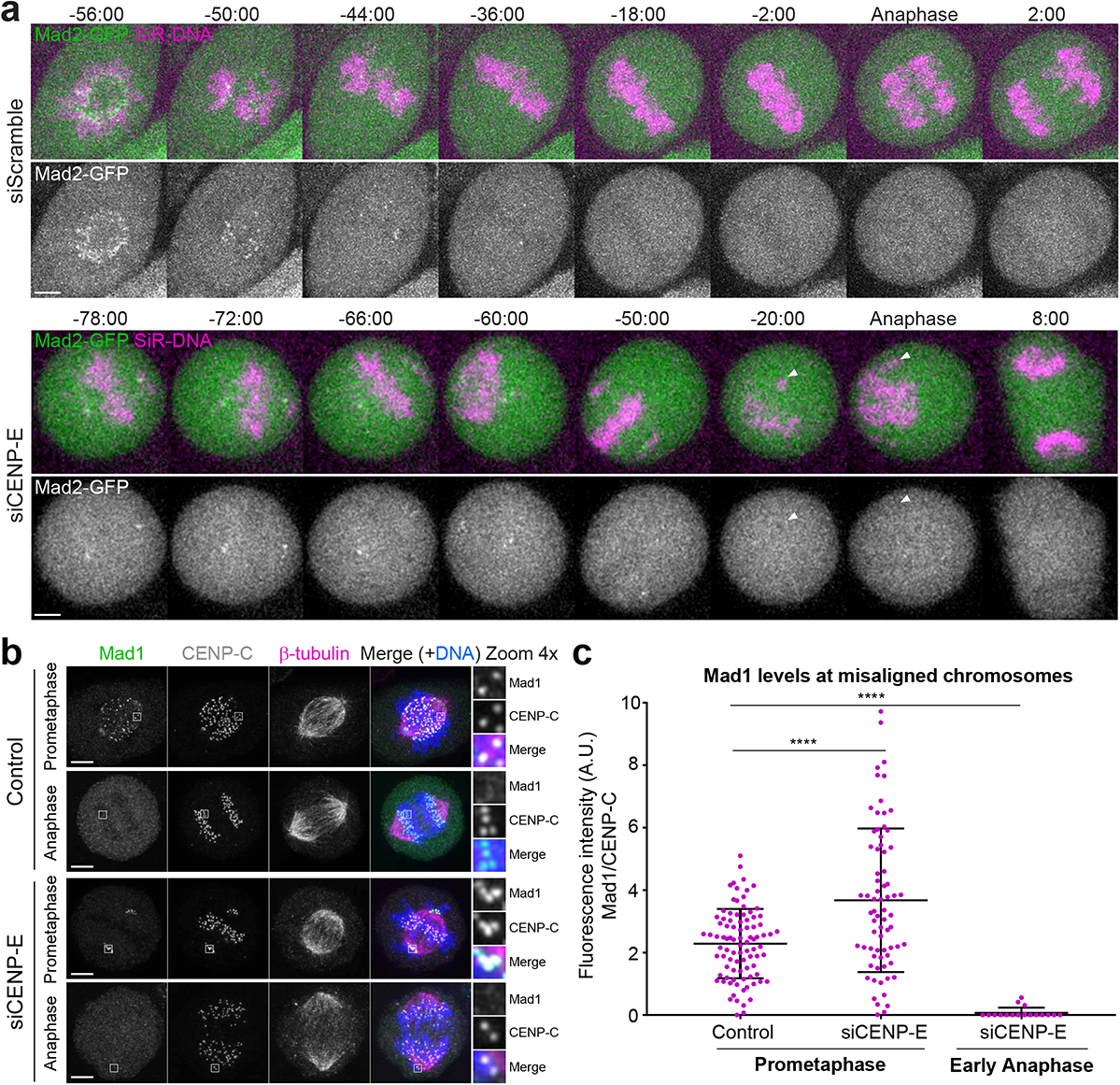
Cells with chronically misaligned chromosomes enter anaphase after satisfying the spindle assembly checkpoint. **a)** Selected time-frames of representative HeLa cells stably expressing Mad2-GFP (green) and chromosomes labeled with SiR-DNA (magenta) in control and after CENP-E depletion. White arrowheads point to a misaligned chromosome during anaphase. Images were acquired every 2 min. Time = min:sec. Time 00:00 = anaphase onset. **b)** Representative immunofluorescence images of HeLa cells stained for DNA (blue), Mad1 (green), CENP-C (white) and β-tubulin (magenta). Insets show higher magnification of selected regions with misaligned chromosomes (grayscale for single channels of Mad1 and CENP-C). Images are maximum intensity projections of deconvolved z-stacks. Scale bar = 5 µm. **c)** Quantification of the fluorescence intensity of Mad1 relative to CENP-C on misaligned chromosomes. Each dot represents an individual kinetochore. Mad1 negative values were considered zero due to the high background fluorescence observed in early anaphase cells. The horizontal line indicates the mean of all quantified kinetochores and the error bars represent the standard deviation from a pool of two independent experiments (mock/prometaphase, 2.286±1.117, n=90 kinetochores, 9 cells; siCENP-E/prometaphase, 3.675±2.298, n=72 kinetochores, 17 cells; siCENP-E/anaphase, 0.06921±0.1653, n=19 kinetochores, 14 cells; ****p≤0.0001 relative to control, analyzed using a Mann-Whitney Test).

To further validate this conclusion, we used time-lapse fluorescence microscopy to quantify the levels and monitor the respective degradation kinetics of Cyclin B1, which was endogenously tagged with the fluorescent protein Venus ^42^ (Figure 5a, b). Consistent with previous reports ^15, 43^, in control HeLa cells Cyclin B1 starts to be steadily degraded few minutes before the onset of anaphase and continues to decline throughout anaphase, becoming undetectable as chromosomes started decondensing in telophase (Figure 5a, b and Movie S7). Similar degradation kinetics were observed in CENP-E-depleted cells that entered anaphase with or without completing chromosome alignment (Figure 5b and Movie S7). We also determined that in this particular set of experiments ∼40% of the CENP-E-depleted anaphase cells formed micronuclei directly from chromosomes that never aligned at the spindle equator (Figure 5c). Lastly, in stark contrast with the very slow Cyclin B1 degradation kinetics over more than 12 hours typical of mitotic slippage/death upon complete microtubule depolymerization with nocodazole (Figure S5a, b) (see also ^40–42^), Cyclin B1 degradation in control- or CENP-E-depleted cells was completed in less than 30 minutes (Figure 5a, b). Taken together, these data indicate that cells with chronically misaligned chromosomes enter anaphase after satisfying the spindle assembly checkpoint and undergoing normal Cyclin B1 degradation.

**Figure 5.**
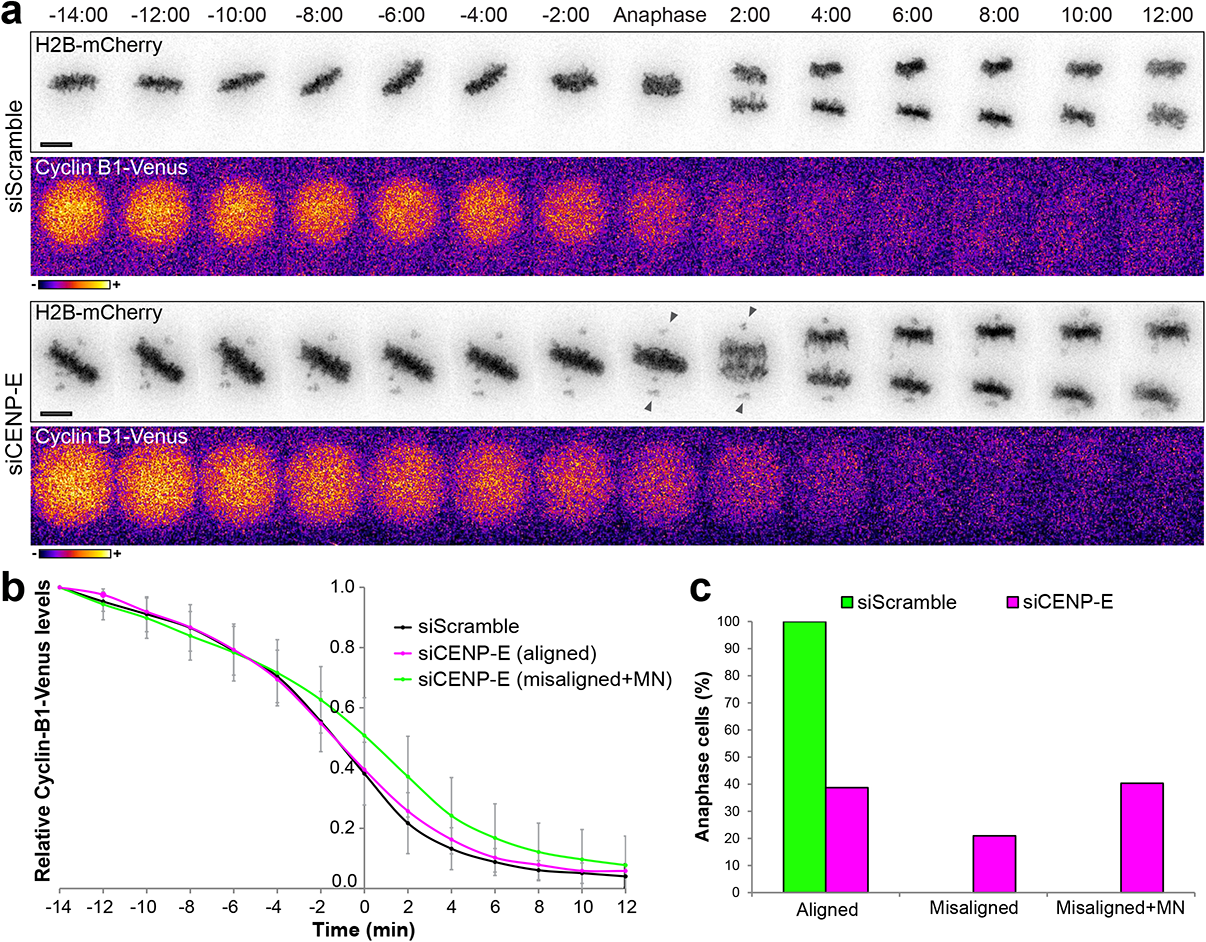
Cells with chronically misaligned chromosomes enter anaphase after undergoing normal Cyclin B1 degradation. **a)** Selected time frames from live-cell microscopy of HeLa cells stably expressing H2B-mCherry and Cyclin B1-Venus in control and CENP-E RNAi. Images were acquired every 2 min. Time = min:sec. Time 00:00 = anaphase onset. Black arrowheads point to misaligned chromosomes during anaphase. **b)** Cyclin B1 degradation curves for control, CENP-E-depleted cells that properly align their chromosomes at the metaphase plate and CENP-E-depleted cells, which exit mitosis with misaligned chromosomes and form micronuclei. The curves represent mean Cyclin B1-Venus fluorescence intensity from all analyzed cells and error bars represent the standard deviation from a pool of two independent experiments (siScramble n=9; siCENP-E (misaligned+micronuclei) n=22; siCENP-E (aligned) n=15). **c)** Frequency of anaphase cells with aligned chromosomes, misaligned chromosomes and misaligned chromosomes that result in micronuclei in control (green bars) and CENP-E-depleted cells (magenta bars).

### Although most micronuclei originate from anaphase lagging chromosomes, chronically misaligned chromosomes are the most faithful predictor of micronuclei formation

Micronuclei are well-established biomarkers of human cancers, cell senescence and genotoxic stress ^44^. The origin of micronuclei has been linked to the presence of lagging chromosomes during anaphase that form due to incorrect merotelic kinetochore-microtubule attachments (when individual kinetochores bind to microtubules oriented to both spindle poles) ^45, 46^. More recently, DNA bridges that persist during anaphase were also implicated in micronuclei formation ^47^. Because in over 100 different molecular perturbations performed, cells frequently entered anaphase with chronically misaligned chromosomes that often missegregate, we sought to compare the relative contributions of lagging and misaligned chromosomes, as well as DNA bridges, to micronuclei formation during HeLa cell division (Figure 6a). To do so, we focused our analysis on a large subset of experimental conditions that are recognized to prevent proper kinetochore-microtubule attachments (Figure 6b). As a rule, and in line with our previous findings ^48^, these conditions led to a substantial increase in the frequency of daughter cells with micronuclei (9.6 ± 9.1%, mean ± s.d., and up to 50% on specific conditions such as KNL1 depletion) when compared to daughter cells treated with a control siRNA (1.6%) (Figure 6b). As expected, most of the resulting micronuclei derived from anaphase lagging chromosomes (62 ± 20%, mean ± s.d. of all conditions) and only very few (4.3 ± 6.4%, mean ± s.d. of all conditions) originated from DNA bridges (Figure 6b). However, we also found that a significant fraction of cells (33 ± 11%, mean ± s.d. of all conditions) formed micronuclei that derived directly from misaligned chromosomes (Figure 6b). Noteworthy, although occurring at much lower frequency, the relative origin of micronuclei in control HeLa cells was identical to that generally observed upon experimental perturbation of kinetochore-microtubule attachments (58%, 10% and 32%, for lagging chromosomes, DNA bridges and misaligned chromosomes, respectively; n=897 cells) (Figure 6b).

**Figure 6.**
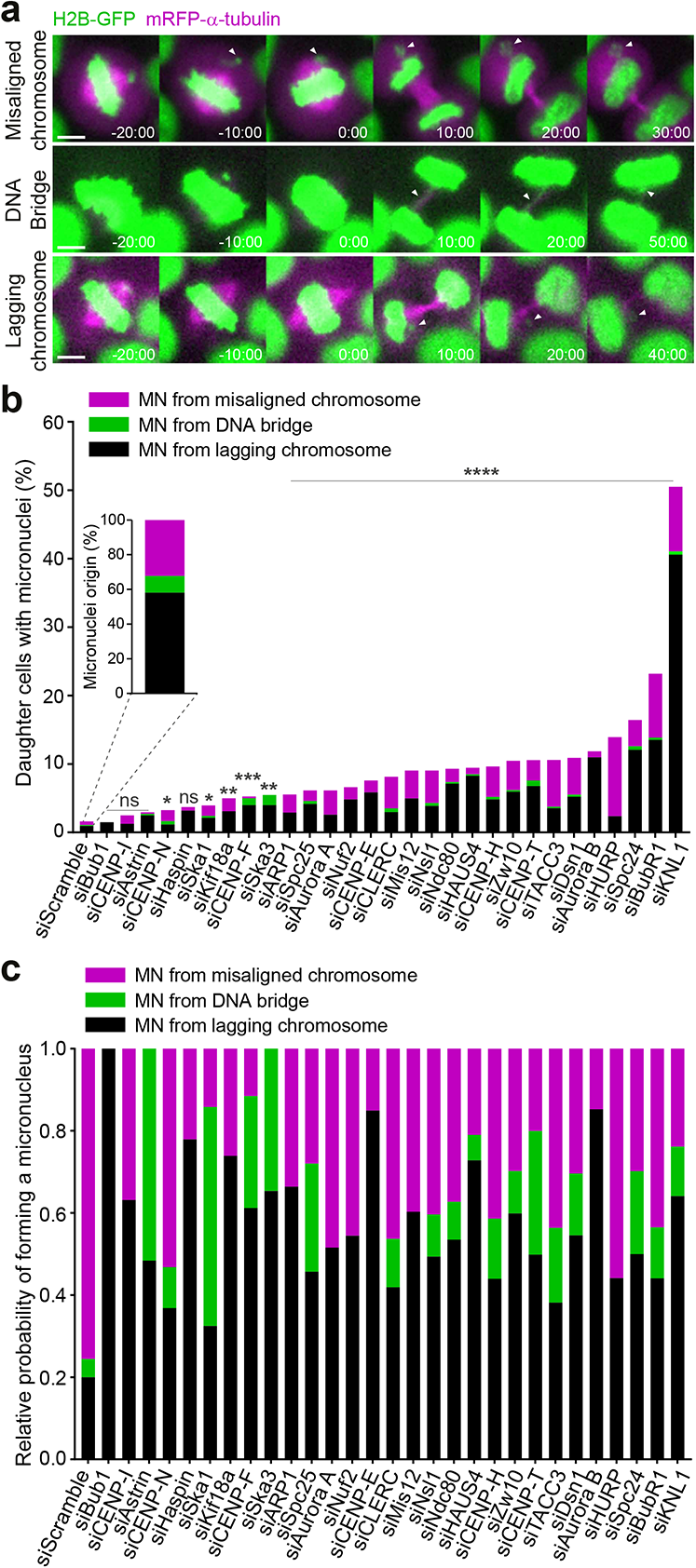
Although most micronuclei originate from anaphase lagging chromosomes, chronically misaligned chromosomes are the most faithful predictor of micronuclei formation. **a)** Examples of time-lapse sequences illustrating possible origins of micronuclei: micronuclei originated from a misaligned chromosome; micronuclei originated from a broken DNA bridge and micronuclei originated from lagging chromosomes. Time = min:sec. Time 00:00 = anaphase onset. White arrowheads track misaligned chromosomes, DNA bridges or lagging chromosomes until they eventually form micronuclei. Pixels were saturated for optimal visualization of misaligned chromosomes, DNA bridges and lagging chromosomes. Scale bar = 5 µm. **b)** Frequency of daughter cells with micronuclei that derived either from lagging chromosomes (black bars), DNA bridges (green bars) or misaligned chromosomes (magenta bars) under the specified conditions [siScramble n=897, siBub1 n=100, siCENP-I n= 98, siAstrin n=199, siCENP-N n= 200, siHaspin n= 100, siSka1 n=146, siKif18a n= 110, siCENP-F n= 200, siSka3 n= 100, siARP1 n=192, siSpc25 n= 222, siAurora A n= 229, siNuf2 n=204, siCENP-E n= 159, siCLERC n=263, siMis12 n= 200, siNsl1 n=200, siNdc80 n=202, siHAUS4 n= 240, siCENP-H n=125, siZw10 n= 187, siCENP-T n=150, siTACC3 n=200, siDsn1 n=459, siAurora B n=287, siHURP n=296, siSpc24 n=182, siBubR1 n= 151, siKNL1 n=194; pool of 2 independent experiments with the exception of the results presented for depletions of Bub1, CENP-I, Kif18a and Ska3 in which only 1 experiment was performed, and CLERC and Dsn1 in which 3 independent experiments were performed; (*p≤0.05, **p≤0.01, ***p≤0.001, ****p≤0.0001, ns corresponds to not significantly different from control; Fisheŕs exact two-tailed test)]. **c)** Relative probability of micronuclei formation from a lagging chromosome (black bars), a DNA bridge (green bars) or a misaligned chromosome (magenta bars) under the specified conditions.

Next, we determined the respective probabilities of micronuclei formation given a specific condition, which can be either a lagging chromosome, a DNA bridge or a misaligned chromosome. Surprisingly, and despite the fact that most micronuclei derive from anaphase lagging chromosomes, we found that in unperturbed HeLa cells treated with a control siRNA, the relative probability of micronuclei formation from a chronically misaligned chromosome clearly outcompeted (0.76) all the other classes, including anaphase lagging chromosomes (0.20) (Figure 6c) (for absolute probabilities see Figure S6a). Although somewhat unexpected, this is not necessarily an oddity: although most people die of cancer or heart diseases, the chances of dying on a plane crash are definitely higher. Interestingly, experimental perturbation of kinetochore-microtubule attachment stability reverted or attenuated the tendency observed in unperturbed cells (Figure 6c and Figure S6a), consistent with a role of stable kinetochore-microtubule attachments in anaphase error correction and micronuclei prevention from lagging chromosomes ^48^. Thus, although the majority of micronuclei originate from anaphase lagging chromosomes, chronically misaligned chromosomes are the most faithful predictor of micronuclei formation during HeLa cell division.

### Micronuclei formation from chronically misaligned chromosomes is a frequent outcome in chromosomally unstable cancer cells, but not in non-transformed cells

Intrigued by the fact that unperturbed HeLa cells sporadically enter anaphase with misaligned chromosomes, which showed the highest risk of forming micronuclei, we set out to investigate the origin of micronuclei that form spontaneously during cell division in unperturbed RPE-1 and U2OS cell lines, two well-established models of near-diploid chromosomally stable (non-transformed) and chromosomally unstable (transformed) human cells, respectively ^49^. To visualize the entire chromosome set and spindle microtubules, these cell lines were engineered to stably express Histone H2B-GFP and mRFP-α-tubulin and were inspected by 4D live-cell spinning-disk confocal microscopy, with 30 sec temporal resolution (Figure 7a, b). In parallel, we promoted chromosome missegregation by performing a monastrol treatment and washout in both cell lines to induce the formation of erroneous kinetochore-microtubule attachments ^50, 51^. Unperturbed RPE-1 cells showed only a residual (0.73%) formation of micronuclei after cell division and none derived from a chronically misaligned chromosome (Figure 7c). As expected, monastrol treatment and washout in RPE-1 cells significantly increased the frequency of daughter cells that resulted in micronuclei (6.5%), most of which (83%) derived from anaphase lagging chromosomes and only in 1 out of 44 recorded cells (corresponding to 17% of the cells that formed micronuclei) derived from a misaligned chromosome (Figure 7c). This scenario was strikingly different even in unperturbed U2OS cells, which formed micronuclei in 7.3% of the cases, of which 52% derived from anaphase lagging chromosomes, 14% from DNA bridges and 34% from chronically misaligned chromosomes (Figure 7c). Monastrol treatment and washout only slightly increased (without statistical significance) the percentage of dividing U2OS cells that formed micronuclei, which in this case derived mostly from anaphase lagging chromosomes (80%), likely due to an increase in merotelic attachments ^50^.

**Figure 7.**
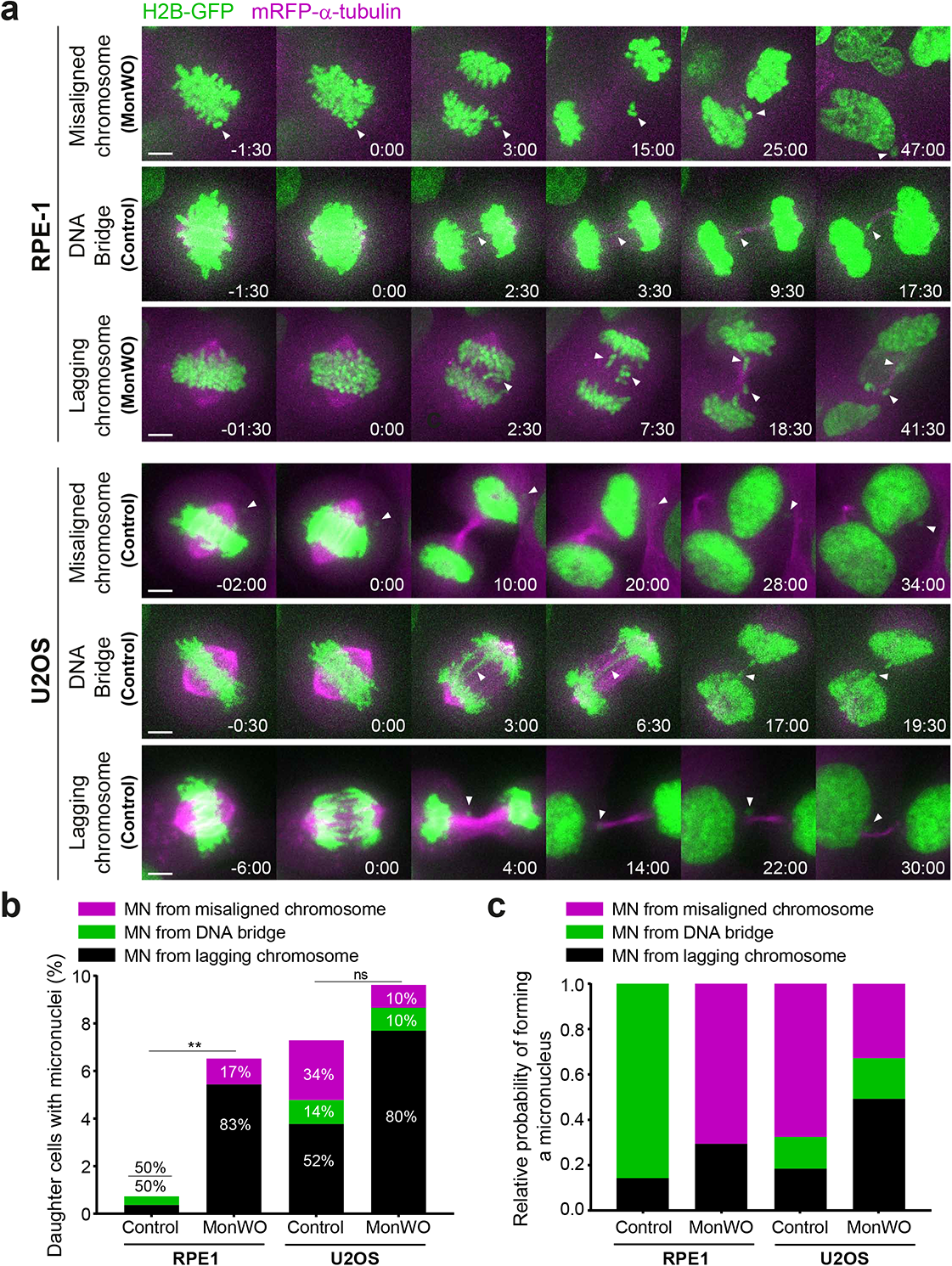
Micronuclei formation from chronically misaligned chromosomes is a frequent outcome in chromosomally unstable cancer cells, but not in non-transformed cells. **a)** Examples of time-lapse sequences illustrating possible origins of micronuclei in RPE-1 and U2OS cells. Time = min:sec. Time 00:00 = anaphase onset. White arrowheads track misaligned chromosomes, DNA bridges or lagging chromosomes until they eventually form micronuclei. Pixels were saturated for optimal visualization of misaligned chromosomes, DNA bridges and lagging chromosomes. Scale bar = 5 µm. **b)** Frequency of RPE-1 and U2OS daughter cells with micronuclei that derived either from lagging chromosomes (black bars), DNA bridges (green bars) or misaligned chromosomes (magenta bars) in control and after monastrol washout. RPE-1 cells: [control n=136, monastrol washout n=44, p=0.0039, analyzed using the Fisheŕs exact two-tailed test]. U2OS cells: [control n=198, monastrol washout n=49, p=0.4150, analyzed using the Fisheŕs exact two-tailed test]. **c)** Relative probability of micronuclei formation from a lagging chromosome (black bars), a DNA bridge (green bars) or misaligned chromosome (magenta bars) in RPE-1 and U2OS cells in control and after monastrol washout.

We next determined the relative probabilities of micronuclei formation from lagging chromosomes, DNA bridges and misaligned chromosomes scored in both RPE-1 and U2OS cells, with and without monastrol treatment and washout (Figure 7d) (for absolute probabilities see Figure S6b). In line with our previous observations in HeLa cells (Figure 6c and Figure S6a), this analysis revealed that chronically misaligned chromosomes have the highest probability of resulting in micronuclei in unperturbed chromosomally unstable U2OS cells (0.68) (Figure 7d and Figure S6b). Interestingly, monastrol treatment and washout reverted this tendency in U2OS cells, once again, likely due to an increase in merotelic attachments (Figure 7d and Figure S6b) ^50^. Most striking, and in sharp contrast to unperturbed HeLa and U2OS cells, unperturbed RPE-1 cells always entered anaphase after completing chromosome alignment and, consequently, no micronuclei from misaligned chromosomes were ever detected in our recordings (Figure 7d and Figure S6b). Nevertheless, in the very few RPE-1 cells that entered anaphase with misaligned chromosomes upon monastrol treatment and washout, chronically misaligned chromosomes showed the highest probability (0.71) among all types of segregation errors of forming micronuclei (Figure 7d and Figure S6b). We conclude that chronically misaligned chromosomes that form spontaneously in unperturbed chromosomally unstable cancer cells, or upon the experimental induction of erroneous kinetochore-microtubule attachments in non-transformed cells, have the highest probability to missegregate and result in micronuclei.

Chromosomally unstable cancer cells are known to have an intrinsic increase in kinetochore-microtubule attachment stability and a poor error correction capacity ^49, 52^. To investigate whether increased kinetochore-microtubule attachment stability in chromosomally unstable cancer cells might allow misaligned chromosomes to satisfy the SAC, we implemented an experimental protocol that first promotes the formation of few misaligned chromosomes after nocodazole treatment and washout (Figure S7a) (see Materials and Methods), followed by a nocodazole shock and quantification of fluorescence intensity from depolymerizing microtubules over time in fixed cells (Figure S7a). Both qualitative and quantitative analyses revealed that, under these experimental conditions, kinetochore microtubules in chromosomally unstable U2OS cells are more resistant to depolymerization induced by a nocodazole shock, when compared to normal near-diploid RPE-1 cells (Figure S7a, b). Measurement of the respective half-life of polymerized tubulin, confirmed ∼2-fold increase in U2OS cells relative to RPE1 cells (Figure S7a, b). These results provide an explanation for the inefficient correction of few misaligned chromosomes that eventually satisfy the SAC in cancer cell models and thus may represent major drivers of chromosomal instability and micronuclei formation in human cancers.

## Discussion

It is currently thought that anaphase lagging chromosomes resulting from erroneous merotelic attachments that satisfy the SAC are major drivers of genomic instability in human cancers (for reviews see ^53, 54^). Although anaphase lagging chromosomes resulting from merotelic attachments rarely missegregate ^55, 56^, they may fail to incorporate into the respective daughter nuclei during cell division and result in the formation of micronuclei. Micronuclei were recently implicated as key intermediates of chromothripsis, a series of massive genomic rearrangements that may drive rapid tumor evolution and account for acquired drug resistance and oncogene activation ^45, 57–60^. We now show that although most micronuclei derive from anaphase lagging chromosomes, simply because these events occur at a very high frequency in chromosomally unstable cancer cells ^55^, chronically misaligned chromosomes that satisfy the SAC often missegregate and have the highest probability to form micronuclei, specifically in human cancer cell models (see graphical abstract). This is consistent with recent high-resolution live-cell studies in both cancer and non-cancer human cells that showed that the vast majority of lagging chromosomes have a transient nature and are corrected during anaphase by an Aurora B-dependent mechanism that prevents micronuclei formation ^48, 61^.

Defects in chromosome alignment are normally avoided by increased Aurora B activity at centromeres of misaligned chromosomes ^27^, as well as by increased Aurora A activity at spindle poles ^5, 62^, causing the destabilization of kinetochore-microtubule attachments and preventing SAC satisfaction. This provides time for misaligned chromosomes to congress to the spindle equator by CENP-E-mediated lateral transport along detyrosinated spindle microtubules ^8, 10^. However, as opposed to normal cells, correction of erroneous attachments underlying chromosome alignment defects (e.g. syntelic attachments in which both kinetochores of a chromosome are oriented towards the same spindle pole) appears to be less robust in cancer cells that also show overly stabilized kinetochore-microtubule attachments ^49, 52^. Interestingly, RPE-1 cells treated with microtubule-targeting drugs at concentrations that stabilize microtubules satisfy the SAC in the presence of misaligned chromosomes, and do so faster under conditions that promote the formation of syntelic attachments ^63, 64^. Together with the fact that normal human near-diploid cells rely on a robust p53-dependent mechanism that limits the proliferation of aneuploid cells ^65^, these data help to explain how spontaneous misaligned chromosomes in cancer cells eventually satisfy the SAC and may represent a direct route to chromosomal stability.

Besides the demonstration of spontaneous formation of micronuclei from misaligned chromosomes in cancer cell models, the present work also unveils a wide range of genetic perturbations that predispose for these events and might account for the underlying chromosomal and genomic instability commonly observed in human cancers. A paradigmatic case is the perturbation of CENP-E function that has been linked to tumorigenesis in vivo ^31^. Previous studies have shown that ∼40% of CENP-E-depleted HeLa cells enter anaphase with misaligned chromosomes ^27, 28^. Fixed cell analysis revealed that these misaligned chromosomes accumulate Mad2, but micronuclei generated from CENP-E-depleted cells did not, suggesting that misaligned chromosomes satisfy the SAC ^27^. Although suggestive, the origin of the scored micronuclei was not determined in these fixed cell experiments, and so it remains possible that the scored micronuclei do not derive directly from misaligned chromosomes (they may alternatively derive from anaphase lagging chromosomes, see Figure 6b,c), and cells with misaligned chromosomes enter anaphase without satisfying the SAC. In fact, previous experiments in fixed CENP-E KO MEFs revealed continued localization of SAC proteins at misaligned chromosomes seen in anaphase cells, suggesting ongoing SAC signaling that was nevertheless insufficient to prevent anaphase onset ^26^. In line with these findings, recent experiments in which CENP-E activity was inhibited in human RPE-1 cells suggest that misaligned chromosomes resulting in micronuclei do so without satisfying the SAC, possibly due to the “ensheathing” of misaligned chromosomes by endomembranes ^66^. Our live-cell imaging of Mad2-GFP upon CENP-E depletion in HeLa cells, supported by quantitative analyses in fixed cells soon after anaphase onset, clarify these discrepancies. Accordingly, we found that Mad1/Mad2 dissociate from kinetochores of chronically misaligned chromosomes in cells that enter anaphase, suggesting SAC satisfaction. Indeed, monitoring fluorescently-tagged endogenous Cyclin B1 by live-cell imaging revealed a normal degradation kinetics in CENP-E-depleted cells that enter anaphase with misaligned chromosomes, when compared with anaphase cells (with or without CENP-E) that complete alignment. This pattern contrasts with that observed upon mitotic slippage, in which mitotic cells that cannot satisfy the SAC exit mitosis with high Mad1/Mad2 levels at kinetochores and after a very slow and prolonged degradation of Cyclin B1 ^41, 64, 67^. Combined, these data provide direct evidence that, at least in certain conditions, cancer cells with chronically misaligned chromosomes may enter anaphase after SAC satisfaction and give rise to micronuclei (see graphical abstract).

Interestingly, our analysis of more than 100 different molecular perturbations indicates that entering anaphase with chronically misaligned chromosomes might be a frequent outcome in cancer cells. Of particular interest, some perturbations, such as CENP-E or Kif18a depletion, were largely compatible with cell viability, despite the high incidence of cells that entered anaphase in the presence of chronically misaligned chromosomes. This contrasts with more drastic scenarios that result from perturbation of end-on kinetochore-microtubule attachments (e.g. depletion of KMN components) that often result in massive chromosome missegregation and cell death. Noteworthy, while the loss of Kif18a, which causes asynchronous segregation of misaligned chromosomes due to loss of interchromosome compaction during anaphase, does not promote chromosomal instability and tumorigenesis ^2, 76^, the loss of CENP-E that typically originates one or few pole-proximal chromosomes directly leads to aneuploidy and the spontaneous formation of lymphomas and lung tumors in aged animals ^26, 31^. Another non-mutually exclusive possibility is that the origin of the resulting micronuclei determines their properties and respective propensity to undergo massive chromosome rearrangements, such as those commonly observed in chromothripsis. For instance, micronuclei derived from segregation errors associated with Kif18a loss of function appear to form stable nuclear envelopes ^76^. However, because misaligned chromosomes that form after perturbation of CENP-E function are brought very close to Aurora A activity at the spindle poles ^5^, this might compromise proper nuclear envelope formation ^77, 78^. In agreement, micronuclei derived from misaligned chromosomes after CENP-E perturbation were recently suggested to activate the cGAS-STING pathway in cancer cells ^79^. Overall, our findings hint for genetically-determined mechanisms underlying micronuclei origin and incite for an in-depth characterization of the properties and fate of micronuclei of different origins, while evaluating their respective potential to drive and/or sustain cell transformation.

## Supplemental Figure Legends

**Figure S1 (related to** Figure 1**). Representative immunoblots of cell lysates to validate the efficiency of depletion of some siRNAs used in the screening.** Protein lysates obtained after RNAi were immunoblotted with an antibody specific for each protein of interest (upper band), except for spc25, spc24 and Nuf2 where anti-Hec1 was used. The bottom band corresponds to anti-GAPDH (siKif2a, siSka2, siBub1, siAurora A, siNdc80, siSpc24, siSpc25, siNuf2, siSeptin-2, siHAUS6, siSurvivin), anti-α-tubulin (siKif4a, siBubR1, siAstrin, siCLASP1, siCLASP2, siCENP-I, siINCENP, siNde1, siAnd-1, siHURP, siDsn1, si4.1r, siCENP-E, siAurora B) and anti-vinculin (siCENP-F, siATRX, siDHC), which were used as loading controls.

**Figure S2 (related to** Figure 1**). Identification of off-targeting effects.** Protein lysates obtained after RNAi treatment were immunoblotted with an antibody specific for each protein of interest (SHP2, GAK, CEP72 and CEP90, upper bands). The bottom band corresponds to anti-GAPDH, which was used as a loading control.

**Figure S3 (related to** Figure 2**). In addition to chromosome alignment defects, some genetic conditions also compromise the maintenance of chromosome alignment at the metaphase plate. a)** Examples of time-lapse sequences illustrating the three main mitotic phenotypes of chromosome alignment maintenance defects observed: 1) cells showed a prolonged delay in chromosome alignment but eventually completed congression, after which chromosomes/chromatids underwent gradual scattering from the metaphase plate; 2) chromosomes aligned normally at the metaphase plate, but then underwent gradual scattering; 3) chromosomes aligned normally at the metaphase plate, followed by spindle pole fragmentation and chromosome scattering. Arrows indicate scattered chromosomes/chromatids in cells that were unable to maintain chromosome alignment at the metaphase plate. Scale bar = 5 µm. Time: h:min, from nuclear envelope breakdown (NEB) to cell death or mitotic exit. **b)** Frequency of cells exhibiting problems in the maintenance of chromosome alignment at the metaphase plate. Only the conditions exhibiting problems in the maintenance of chromosome alignment were included. At least 2 independent experiments were analyzed. The total number of cells analyzed for each condition is indicated in Table S1. (*p≤0.05, **p≤0.01, ***p≤0.001, ****p≤0.0001, ns corresponds to not significantly different from control, Fisheŕs exact two-tailed test).

**Figure S4 (related to** Figure 3**). Mitotic timing upon gene-specific RNAi-mediated depletion**. HeLa cells stably expressing H2B-GFP and α-tubulin-mRFP were acquired every 10 minutes. Mitotic duration was determined by measuring the time between nuclear envelope breakdown (NEBD) and anaphase onset (AO), shown in minutes. Each data point corresponds to one cell, the magenta rectangle represents the mean value, and the error bars represent the standard deviation from a pool of at least 2 independent experiments. The difference between mean values of each RNAi condition was statistically significant from the control mean values (*p≤0.05, **p≤0.01, ***p≤0.001, ****p≤0.0001, ns corresponds to not significantly different from control, Mann-Whitney test)

**Figure S5 (related to** Figure 5**). Cyclin B1-venus degradation upon complete microtubule depolymerization with nocodazole. a)** Selected time frames from phase contrast and fluorescence microscopy of Cyclin B1-Venus HeLa cells treated with nocodazole with or without MG132. Images were acquired every 15 min. Scale bar = 5 µm. Time = h:min. **b)** Cyclin B1 degradation curves for nocodazole treated Cyclin B1-Venus HeLa cells in the presence or absence of MG132. Fluorescence intensities were normalized to the levels at time = 0. The curves depict mean Cyclin B1-Venus fluorescence intensity from all analyzed cells per condition (nocodazole n=12; nocodazole + MG132 n=10), and error bars represent the standard deviation. Note that acquisition in the presence of MG132 was terminated earlier relative to acquisition without MG132 due to cell death.

**Figure S6 (related to** Figure 6 **and** Figure 7**). Absolute probabilities of forming a micronucleus from misaligned chromosomes, DNA bridges and lagging chromosomes in the different cell lines used. a)** Absolute probabilities of forming a micronucleus of different origins in unperturbed HeLa cells and after molecular perturbations that weaken kinetochore-microtubule attachments. [siScramble n=897, siBub1 n=100, siCENP-I n= 98, siAstrin n=199, siCENP-N n= 200, siHaspin n= 100, siSka1 n=146, siKif18a n= 110, siCENP-F n= 200, siSka3 n= 100, siARP1 n=192, siSpc25 n= 222, siAurora A n= 229, siNuf2 n=204, siCENP-E n= 159, siCLERC n=263, siMis12 n= 200, siNsl1 n=200, siNdc80 n=202, siHAUS4 n= 240, siCENP-H n=125, siZw10 n= 187, siCENP-T n=150, siTACC3 n=200, siDsn1 n=459, siAurora B n=287, siHURP n=296, siSpc24 n=182, siBubR1 n= 151, siKNL1 n=194; pool of 2 independent experiments with the exception of the results presented for depletions of Bub1, CENP-I, Kif18a and Ska3 in which only 1 experiment was performed, and CLERC and Dsn1 in which 3 independent experiments were performed. **b)** Absolute probabilities of forming a micronucleus of different origins in RPE-1 and U2OS cells, with and without Monastrol treatment and washout. RPE-1 cells: [control n=136, monastrol washout n=44]. U2OS cells: [control n=198, monastrol washout n=49].

**Figure S7 (related to** Figure 7**). Kinetochore-microtubule attachments associated with misaligned chromosomes are more stable in cancer cells. a)** Representative immunofluorescence images of RPE-1 and U2OS cells stained for DNA (green) and α-tubulin (magenta). RPE-1 and U2OS cells upon nocodazole treatment and washout to generate misaligned chromosomes were processed for immunofluorescence microscopy after a subsequent nocodazole shock 5, 15 and 30 min after drug addition. Representative immunofluorescence images of the mitotic spindle at each stage are shown. Images are maximum intensity projections of deconvolved z-stacks. Scale bar = 5 µm. **b)** Normalized α-tubulin fluorescence intensity at indicated time points in RPE-1 and U2OS cells after nocodazole shock. Data represent mean ± s.d., U2OS n=22 cells, RPE-1 n=22 cells, from 2 independent experiments. Whole lines show single exponential fitting curve (**p≤0.01, extra sum-of-squares F test).

## Supplemental Movie Legends

**Movie S1 (related to** Figure 2). Time-lapse recording of control siScramble-depleted HeLa cells stably expressing H2B-GFP and α-tubulin-mRFP to visualize chromosomes (green) and microtubules (magenta). Time = h:min. Only the first 24h are shown. (For better visualization, open file in ImageJ and zoom in on individual cells)

**Movie S2 (related to** Figure 2 **and S3).** Time-lapse recording of HeLa cells stably expressing H2B-GFP and α-tubulin-mRFP to visualize chromosomes (green) and microtubules (magenta) after siRNA depletion of Augmin, CLASPs and Ska complex proteins. Time = h:min. Only the first 24h are shown. (For better visualization, open file in ImageJ and zoom in on individual cells)

**Movie S3 (related to** Figure 2 and 3). Time-lapse recording of HeLa cells stably expressing H2B-GFP and α-tubulin-mRFP to visualize chromosomes (green) and microtubules (magenta) after siRNA depletion of CENP-E, TRAMM and Dynein Heavy Chain (DHC). Time = h:min. Only the first 24h are shown. (For better visualization, open file in ImageJ and zoom in on individual cells)

**Movie S4 (related to** Figure 2 **and**3). Time-lapse recording of HeLa cells stably expressing H2B-GFP and α-tubulin-mRFP to visualize chromosomes (green) and microtubules (magenta) after siRNA depletion of KNL1, Mis12 and Ndc80/Hec1. Time = h:min. Only the first 24h are shown. (For better visualization, open file in ImageJ and zoom in on individual cells)

**Movie S5 (related to** Figure 3). Time-lapse recording of HeLa cells stably expressing H2B-GFP and α-tubulin-mRFP to visualize chromosomes (green) and microtubules (magenta) after siRNA depletion of CENP-E and Kif18a. Time = h:min. Only the first 24h are shown. (For better visualization, open file in ImageJ and zoom in on individual cells)

**Movie S6 (related to** Figure 4). Time-lapse recording of HeLa cells stably expressing Mad2-GFP (green) and SiR-DNA to visualize chromosomes (magenta) after control siScramble and CENP-E depletion. Corresponding inverted grayscale panels show Mad2-GFP signal alone. Black arrows indicate Mad2-GFP associated at kinetochores of pole-proximal chromosomes after CENP-E depletion. Time = h:min.

**Movie S7 (related to** Figure 5). Time-lapse recording of HeLa cells stably expressing Cyclin B1-venus (Fire lookup table) and H2B-mCherry to visualize chromosomes (inverted grayscale) after control siScramble and CENP-E depletion. Time = h:min.

## Materials and Methods

### Cell culture and reagents

All cell lines were cultured at 37°C in 5% CO_2_ atmosphere in Dulbeccós modified medium (DMEM, Gibco ^TM^, Thermofisher) containing 10% fetal bovine serum (FBS, Gibco ^TM^, Thermofisher). HeLa H2B-GFP/mRFP-α-tubulin and RPE-1 H2B-eGFP/mCherry-α-tubulin cells were generated by lentiviral transduction. U2OS parental and H2B-eGFP/mCherry-α-tubulin were kindly provided by S. Geley (Innsbruck Medical University, Innsbruck, Austria). hTERT-RPE-1 (RPE-1) parental (ATCC® CRL-400^TM^) was kindly provided by Ben Black (U. Pennsylvania, PA, USA). HeLa Mad2-GFP cells were previously described ^80^. Cyclin B1-Venus expressing HeLa cells were kindly provided by J. Pines (Cancer Research Institute, London, UK). Microtubule depolymerization was induced by nocodazole (Sigma-Aldrich) at 1 µM. To inhibit the proteasome, induce a metaphase arrest, and prevent exit due to a compromised SAC, cells were treated with 5 µM MG132 (EMD Millipore). To promote chromosome missegregation in U2OS and RPE-1 cells, a monastrol washout assay was performed. Briefly, cells were incubated during 8-10 h with 100 µM monastrol. After this period, monastrol was washed twice with warm PBS followed by washing with warm fresh medium and entry in anaphase was monitored under the microscope.

### High-content live-cell imaging RNAi screen

All siRNA sequences used were either a commercial predesigned siRNA from Sigma-Aldrich (MISSION siRNA) or were previously validated by other published studies (see Table S2). For each protein, depletion efficiency was first optimized after preliminary phenotypic analysis between 24-96h upon siRNA transfection (for specific conditions see Table S1) and confirmed by western blotting whenever antibodies against specific proteins were available (Figure S1). For few proteins whose role in chromosome congression remains unclear at the mechanistic level, a second siRNA was used to rule-out possible off-targeting effects. This led to the identification of four proteins (Shp2, GAK, CEP72 and CEP90), where no discernable congression phenotype was observed with the second siRNA, despite a clear reduction in protein levels with both siRNA sequences (Figure S2), suggesting that previously published observations were due to off-targeting effects ^81–84^. Treatment with scramble siRNA was undistinguishable from mock transfection (Lipofectamine only) and was therefore used as a negative control throughout the manuscript. A total of 120 conditions were analyzed in this study. For high-content live-cell imaging, Hela cells stably expressing GFP-H2B/mRFP-α-tubulin were plated onto 96-well plate in DMEM supplemented with 5% FBS and after 1 h transfected with siRNA oligonucleotides (Table S2) at a final concentration of 50 nM. Transfections were performed using Lipofectamine RNAiMAX in Opti-MEM medium (both from Thermo Fisher Scientific) according to the manufacturer’s instructions. Transfection medium was replaced with complete medium after 6 h. For time-lapse microscopy acquisition, cell culture medium was changed to DMEM without phenol red supplemented with 10% FBS 6-12 h before acquisition. Cells were imaged for 72h in an IN CELL Analyzer 2000 microscope (GE Healthcare, Chicago, IL, USA) equipped with temperature and CO_2_ controller, using a Nikon 20x/0.45 NA Plan Fluor objective according to manufacturer instructions. Single planes were acquired every 10 min for approximately 72 h. Images were processed using ImageJ software. Long-term recordings of HeLa Cyclin B1-venus treated with nocodazole and MG132 were also performed under similar condition using in the same high-content microscopy system, imaged every 15 min for 13 h.

### Other RNAi experiments

For high-resolution live cell imaging and immunofluorescence analysis of CENP-E depletion (siCENP-E), cells were plated at 50-60% confluence onto 22 x 22 mm No. 1.5 glass coverslips in DMEM supplemented with 5% of FBS. RNAi transfection was performed using Lipofectamine ^TM^ RNAiMAX reagent (Thermofisher) with 20 nM of siRNA against human CENP-E (see siRNA sequence in table S2), diluted in serum-free media (Opti-MEM^TM^, Thermofisher). Depletion of CENP-E was maximal at 24 h after siRNA transfection and all of the analysis was performed at 24 h.

### High-resolution time-lapse microscopy

For high-resolution time-lapse microscopy, cells were plated onto 22 x 22 mm No. 1.5 glass coverslips (Corning) and cell culture medium was changed to phenol-red-free DMEM CO_2_-independent medium (Invitrogen) supplemented with 10% FBS 6-12 h before mounting. Coverslips were mounted onto 35-mm magnetic chambers (14 mm, no. 1.5, MaTek corporation) immediately before imaging. Time-lapse imaging was performed in a heated chamber (37°C) using a 100x oil-immersion 1.40 NA Plan-Apochromatic objective mounted on an inverted microscope (Eclipse TE2000U; Nikon) equipped with a CSU-X1 spinning-disk confocal head (Yokogawa Corporation of America) controlled by NIS-Elements software and with three laser lines (488nm, 561nm, and 647 nm). Images were detected with a iXonEM+ EM-CCD camera (Andor Technology). Images of U2OS H2B/Tub and RPE H2B/Tub were collected every 2 minutes or 30 seconds: 9 x 2 μm z-stacks spanning a total volume of 16 μm. For imaging of HeLa Mad2-GFP and HeLa cyclin-B1-venus/H2B-mCherry eleven 1-μm-separated z-planes covering the entire volume of the mitotic spindle were collected every 2 min. All displayed images represent maximum-intensity projections of Z-stacks, analysed with the open source image analysis software ImageJ.

### Immunofluorescence microscopy

For immunofluorescence processing, cells were fixed with 4% Paraformaldehyde (Electron Microscopy Sciences) for 10 min followed by extraction with 0.3% Triton X-100 in PBS (Sigma-Aldrich) for 10 min. After blocking with 10% FBS in PBS with 0.1% Triton X-100, all primary antibodies were incubated at 4°C overnight. Then, the cells were washed with PBS containing 0.1% Triton X-100 and incubated with the respective secondary antibodies for 1 h at room temperature. Primary antibodies used were: mouse anti-Mad1 (1:500; Merck Millipore); mouse anti α-tubulin (1:2000; Sigma); rabbit anti-β-tubulin (1:2000; Abcam); anti-guinea pig CENP-C (1:1000; MBL International). Secondary antibodies used were Alexa Fluor 488, Alexa Fluor 568 and Alexa Fluor 647 (1:1500; Themofisher). DNA was counterstained with 1 μg/mL DAPI (4’,6’-diamino-2-fenil-indol; Sigma-Aldrich) and mounted onto glass slides with 20 mM Tris pH8, 0.5 N-propyl gallate and 90% glycerol. Images were acquired using an AxioImager Z1 (63x, Plan oil differential interference contract objective lens, 1.46 NA; from Carl Zeiss), coupled with a CCD camera (ORCA-R2; Hamamatsu Photonics) and the Zen software (Carl Zeiss). Blind deconvolution of 3D image datasets was performed using Autoquant X software (Media Cybernetics).

### Quantification of mitotic errors

Mitotic errors were tracked and quantified manually through the assessment of H2B localization in single plane images. Mitotic errors were divided into 3 main classes: lagging chromosomes, DNA bridges or misaligned chromosomes and these were discriminated according to location and morphology associated with H2B localization. Lagging chromosomes retained normal DNA condensation and emerged at different stages during anaphase. Any H2B-positive material between the two chromosomes masses, but distinguishably separated from them, was counted as lagging chromosomes. DNA bridges were characterized by stretches of DNA that connected both daughter nuclei and often displayed aberrant DNA condensation as judged by H2B localization. Misaligned chromosomes were characterized by any H2B-positive material which remains near the spindle pole. To determine micronuclei origin, fully formed micronuclei were backtracked to reveal whether these originated from lagging chromosomes, DNA bridges or misaligned chromosomes. The absolute probability of micronucleus formation from a lagging chromosome was determined by a ratio between the number of daughter cells with micronuclei derived from lagging chromosomes and the cells with lagging chromosomes. The absolute probability of micronucleus formation from a DNA bridge was determined by a ratio between the number of daughter cells with micronuclei derived from DNA bridges and the cells with DNA bridges. The absolute probability of micronucleus formation from a misaligned chromosome was determined by a ratio between the number of daughter cells with micronuclei derived from misaligned chromosomes and the cells that exit mitosis with a misaligned chromosome. For the relative probabilities, the sum of the 3 independent absolute probability values was normalized to 1.

### Quantitative image analysis

For quantification of Mad1 fluorescence intensity, images were analysed using ImageJ. Briefly, individual kinetochores were identified by CENP-C staining and marked by a region of interest (ROI). The average fluorescence intensity of signals of Mad1 at kinetochores was measured on the focused z plan. The background signal was measured within a neighbouring region and was subtracted from the measured fluorescence intensity the region of interest. Fluorescence intensity measurements were normalized to the CENP-C signals. Mad1 negative values were considered zero, since resulted from the high background fluorescence observed in early anaphase cells. Approximately 90 kinetochore pairs from 9 cells were analysed for control prometaphase cells, 72 kinetochore pairs from 14 cells for prometaphase in CENP-E depleted cells and 19 kinetochore pairs from 14 cells for early anaphase in CENP-E depleted cells. The fluorescence levels of Cyclin B1 in HeLa cells treated with nocodazole were measured using the IN Cell Developer Toolbox software (GE Healthcare). After background subtraction, fluorescence intensities were normalized to the level at time = 0 and represented as a function of time. For the quantifications of cyclin B1 levels in siScramble and siCENP-E HeLa cells, images were analysed using ImageJ. A small square region of interest (ROI) was defined, and cyclin B1 fluorescence intensity measured, throughout time in the cell. The same ROI was used to measure the background outside the region of interest. All fluorescence intensity values were then background corrected and the values were normalized to the highest Cyclin B1 level. The microtubule depolymerization rate after nocodazole treatment in U2OS and RPE1 cells was determined by the proportion of total and soluble α-tubulin levels. The total α -tubulin intensity was measured by drawing a larger oval shaped region of interest (ROI) contained the entire cell in sum-projected images (ImageJ). The soluble α -tubulin levels were determined by drawing five smaller oval shaped ROI outside the chromosome region and the average of these values were calculated in sum-projected images. The fluorescence intensities were normalized to the level at time = 0 and represented as a function of time.

### Western Blotting

Cell extracts were collected after trypsinization and centrifuged at 1200 rpm for 5 min, washed and re-suspended in Lysis Buffer (NP-40, 20 nM HEPES/KOH pH 7.9; 1 mM EDTA pH 8; 1 mM EGTA; 150 mM NaCl; 0.5% NP40; 10% glycerol, 1:50 protease inhibitor; 1:100 Phenylmethylsulfonyl fluoride). The samples were then flash frozen in liquid nitrogen and kept on ice for 30 min. After centrifugation at 14000 rpm for 20 min at 4°C the supernatant was collected and protein concentration determined by the Bradford protein assay (Bio-Rad). Fifty micrograms of total extract were then loaded in SDS-polyacrylamide gels and transferred onto nitrocellulose membranes for western blot analysis. The membranes were blocked with 5% milk in TBS with 0.1% Tween-20 (TBS-T) at room temperature during 1 h, and all primary antibodies were incubated at 4°C overnight. After three washes in TBS-T the membranes were incubated with the secondary antibody for 1 h at room temperature. The membranes were washed in the same conditions than previously and the detection was performed with Clarity Western ECL Substrate (Bio-Rad). The following antibodies were used for western blot: mouse anti-Hec1 (9GA) (1:1000; Abcam), mouse anti-Dsn1 (1:500; a gift from A. Musacchio), mouse anti-CENP-E (1:500; Santa Cruz Biotechnology), mouse anti-Aurora B (1:1000; BD Bioscience), mouse anti-ATRX (1:1000; Santa Cruz Biotechnology), mouse anti-CENP-F (1:1000; BD Biosciences), rabbit anti-CEP72 (1:1000; Novus Biologicals), mouse anti-GAK (1:500; R&D Systems), rabbit anti-WDHD1/And-1 (1:1000; Novus Biologicals), rabbit anti-Aurora-A (1:1000; Novus Biologicals), rabbit anti-HURP (1:200, a gift from Patrick Meraldi), mouse anti-INCENP (1:500; Santa Cruz Biotechnology), rabbit anti-LRRCC1/CLERC (1:1000; Abcam), mouse anti-Sgo-1 (F-8) (1:1000; Santa Cruz Biotechnology), rabbit anti-DHC (1:500; ThermoFisher Scientific), mouse anti-Ned1 (:500; Abnova), sheep anti-Bub1 (1:1000; a gift from Stephen Taylor); rabbit anti-Septin-2 (1:500; Novus Biologicals), rabbit anti-CEP90/PIBF1 (1:1000; Novus Biologicals), mouse anti-Ska2 (1:1000; Santa Cruz Biotechnology), mouse anti-4.1r (B-11) (1:1000; Santa Cruz Biotechnology), rabbit anti-Astrin (N-terminal) (1:1000; a gift from Duane Compton), rabbit anti-Kif4a (1:1000; ThermoFisher Scientific), rat anti-CLASP1 (1:50; Maffini et al 2009), rat anti-CLASP2 (1:50; Maffini et al 2009), rabbit anti-BubR1 (1:500; Abcam), mouse anti-Nde1 (1:1000; Abnova), rabbit anti-SHP2 (1:1000; Abcam), rabbit anti-survivin (1:1000; Novus Biologicals), mouse anti-GAPDH (1:40000; Proteintech), rabbit anti-vinculin (1:1000;ThermoFisher Sientific mouse anti-α-tubulin (clone B-512; 1:5000; Sigma-Aldrich) were used as primary antibodies, and anti-mouse-HRP, anti-rabbit-HRP, anti-sheep-HRP and anti-rat-HRP were used as secondary antibodies (1:5000; Jackson ImmunoResearch Laboratories, Inc.).

### Statistical analysis

All results presented in this manuscript were obtained from pooling data from at least 2 independent experiments unless otherwise stated. Sample sizes and statistical tests used for each experiment are indicated in the respective figure legends. Quantifications of mitotic errors (i.e. cell death and micronuclei) were analyzed using the Fisher’s exact two-tailed test. Correlations were calculated using two-tailed Pearsońs correlation coefficients. When only two experimental groups were compared, we used either a parametric t test or a nonparametric Mann-Whitney test. Distribution normalities were assessed using the D’Agostino-Pearson omnibus test. For the comparison of the single exponential fitting curve extra sum-of square F test was used. For each graph, where applicable, ns= non-significant, *p≤0.05, **p≤0.01 ***p≤0.001 and ****p≤0.0001, unless stated otherwise. In all plots error bars represent standard deviation. All statistical analysis was performed using GraphPad Prism V7 (GraphPad Software, Inc).

### Data availability

A public website where high-resolution time-lapse movie galleries, phenotypic analysis and siRNA sequences for all genes studied will be freely available upon publication.

## Acknowledgments

We thank André Maia and the BioSciences Screening Facility at i3S for technical assistance during this project and members of the Maiato lab for the critical reading of the manuscript. M.G. and M.N.C. are recipients of PhD studentships from Fundação para a Ciência e a Tecnologia of Portugal (SFRH/BD/130938/2017 and SFRH/BD/117063/2016, respectively). This work was funded by the European Research Council (ERC) consolidator grant CODECHECK, under the European Union’s Horizon 2020 research and innovation programme (grant agreement 681443), Fundação para a Ciência e a Tecnologia of Portugal (PTDC/MED-ONC/3479/2020), and a La Caixa Health Research Grant (LCF/PR/HR21/52410025).

## Author contributions

Conceptualization, Supervision, Project Administration and Funding acquisition (HM); Methodology (AMG, BO, MNC, FDS, CF); Investigation, Formal Analysis and Validation (AMG, BO, MNC, FDS, CF, HM); Visualization (AMG, BO, MNC, FDS, CF, HM); Writing – Original Draft (AMG, HM); Writing – Review and Editing (AMG, BO, MNC, CF, HM).

## Declaration of interests

Bernardo Orr declares that he is a consultant specialist at Volastra Therapeutics.

## Notes

### Competing Interest Statement

The authors have declared no competing interest.

