## Supplementary figures and images for "Misaligned chromosomes that satisfy the spindle assembly checkpoint are a strong predictor of micronuclei formation in dividing cancer cells"

### Figure S1

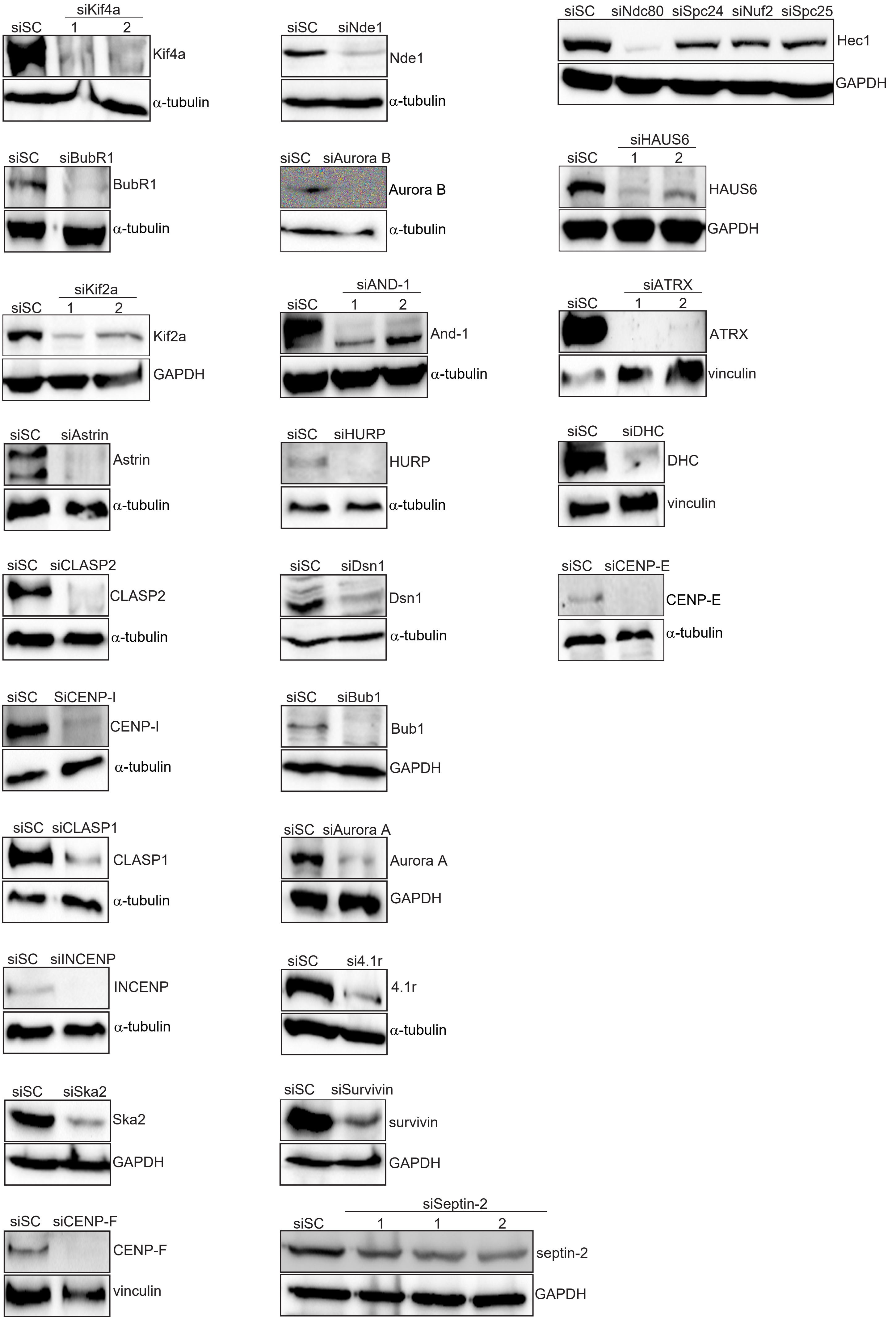


**Gomes et al - Figure S1**

### Figure S2

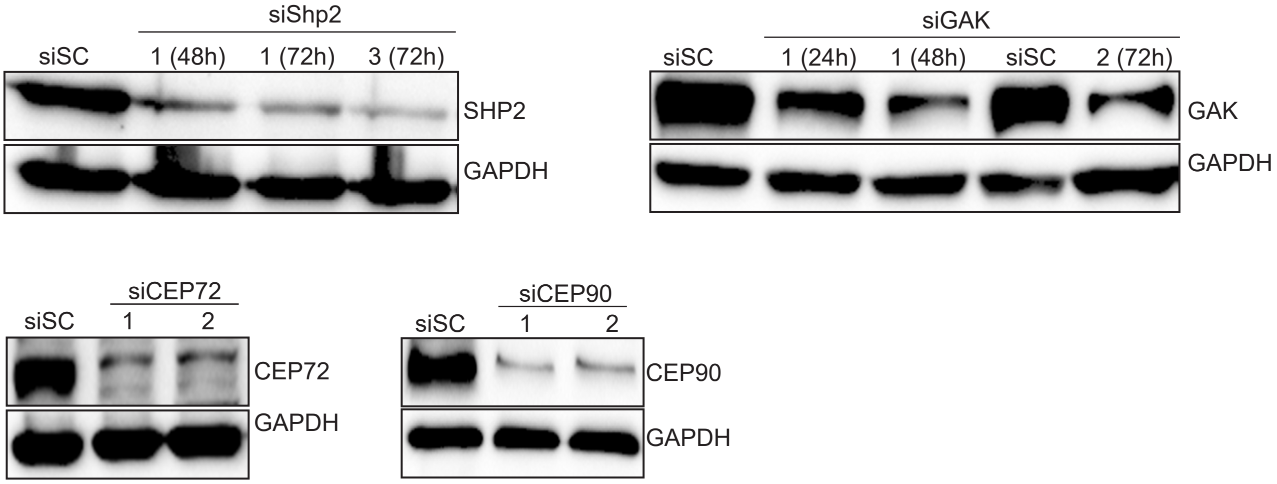


**Gomes et al - Figure S2**

### Figure S3

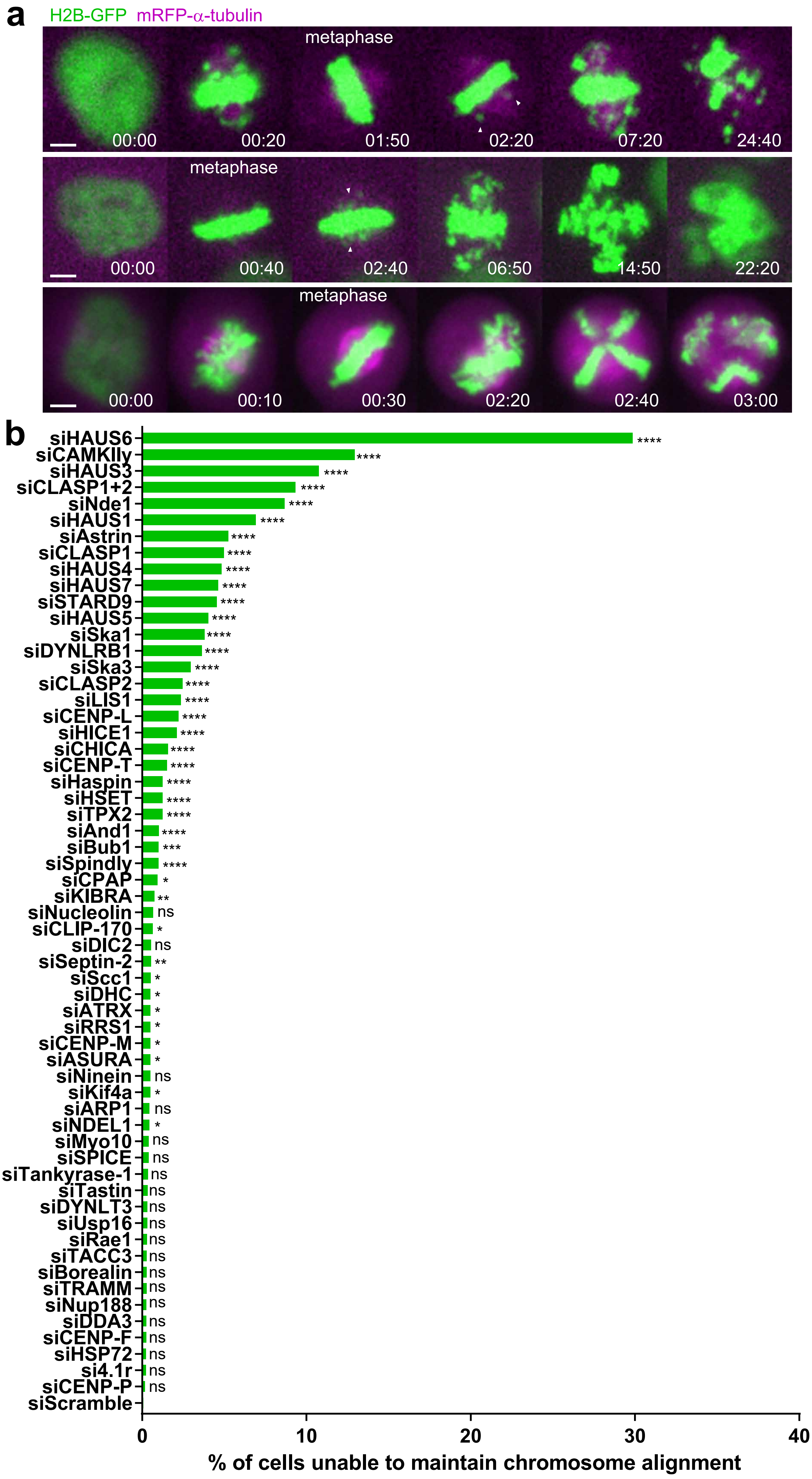


Gomes et al - Figure S3

### Figure S4

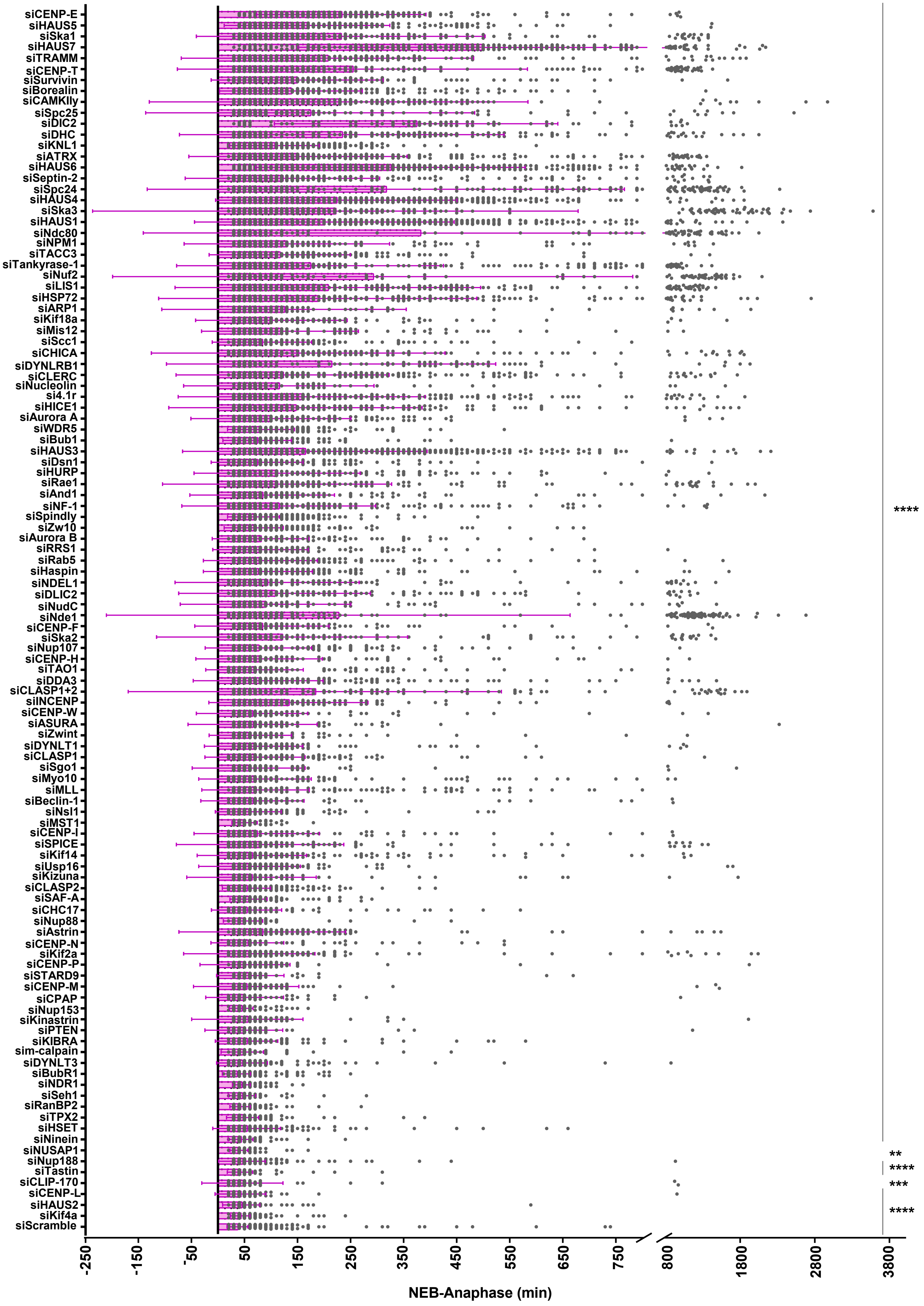


**Gomes et al - Figure S4**

### Figure S5

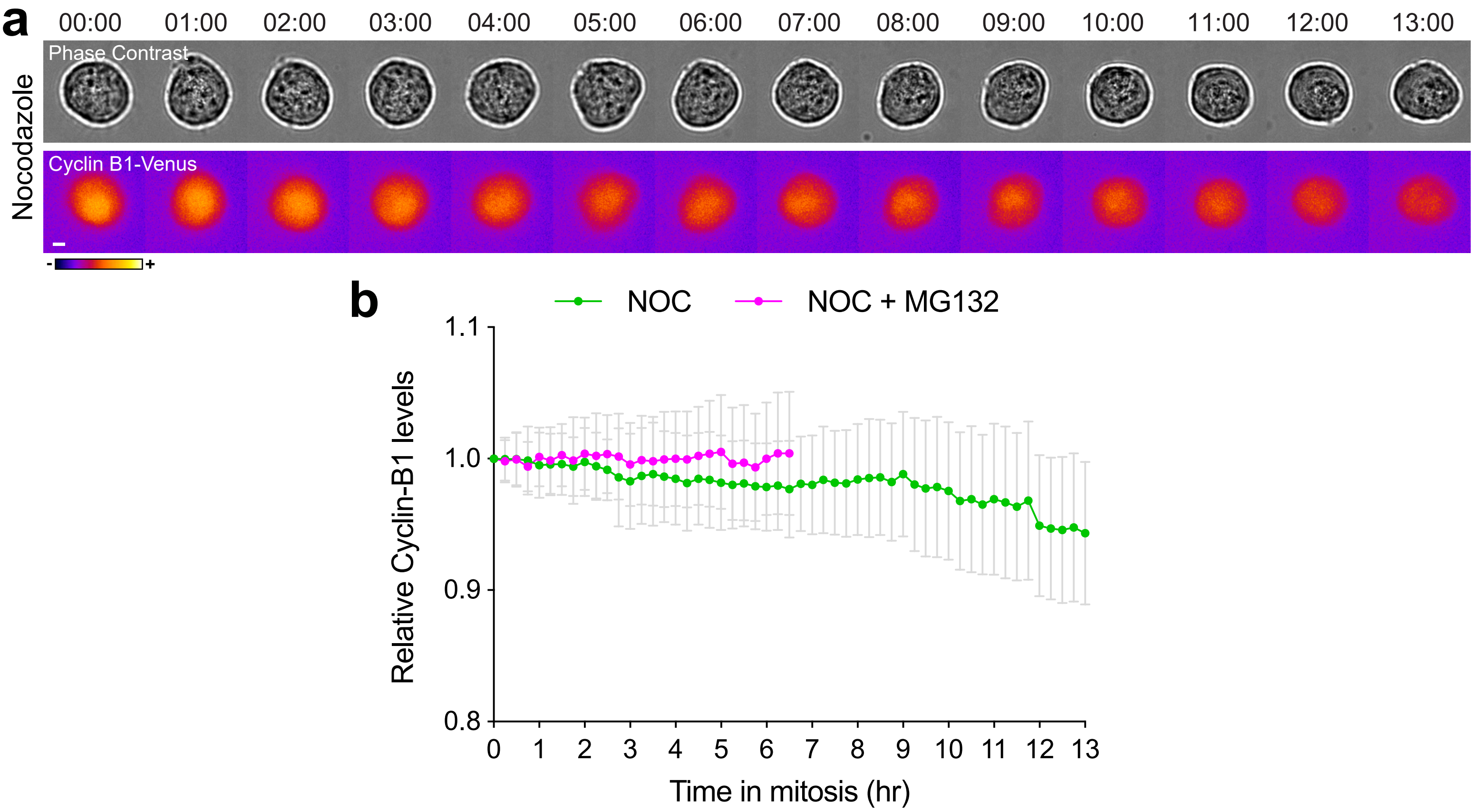


**Gomes et al - Figure S5**

### Figure S6

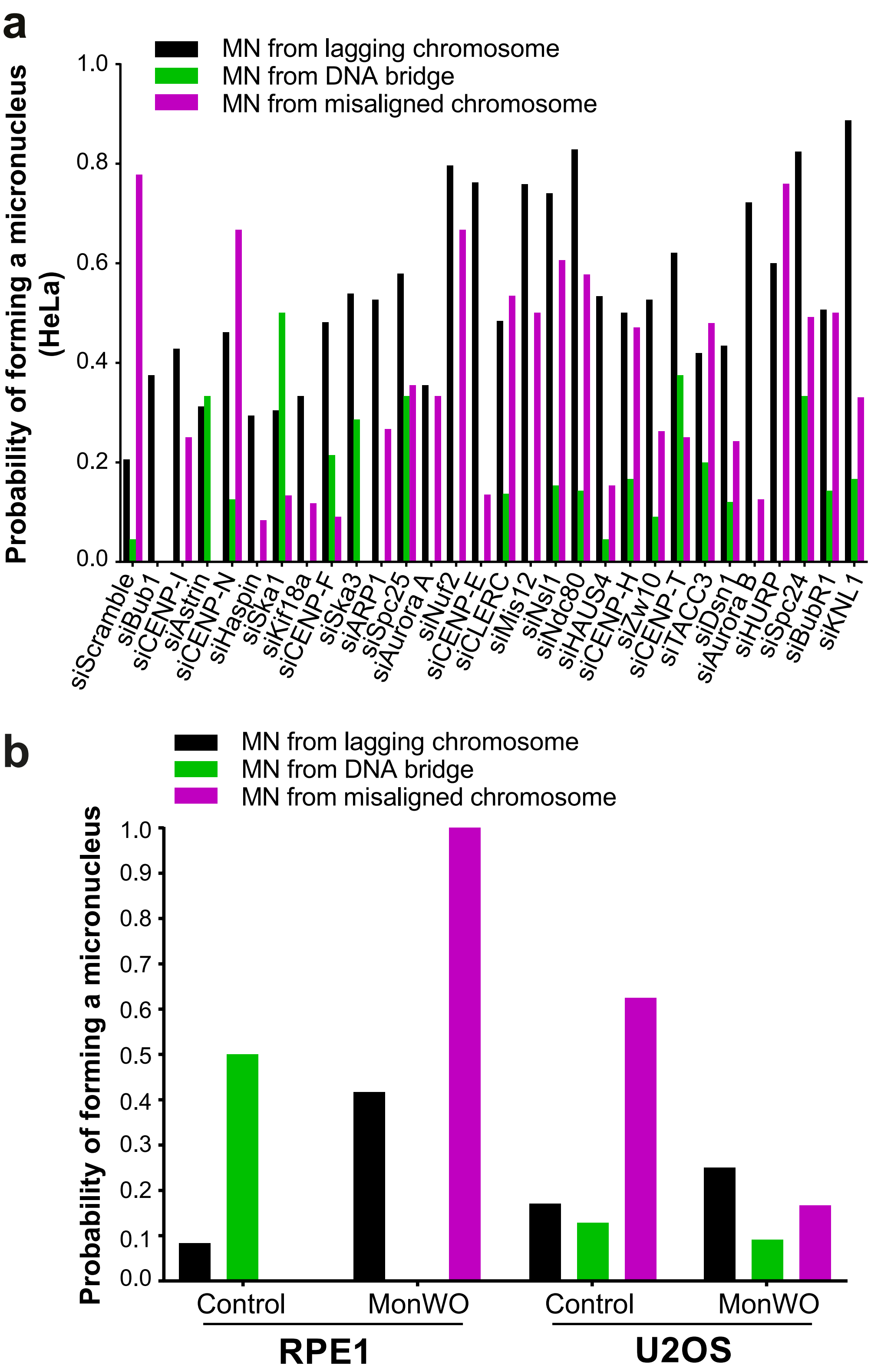


Gomes et al - Figure S6

### Figure S7

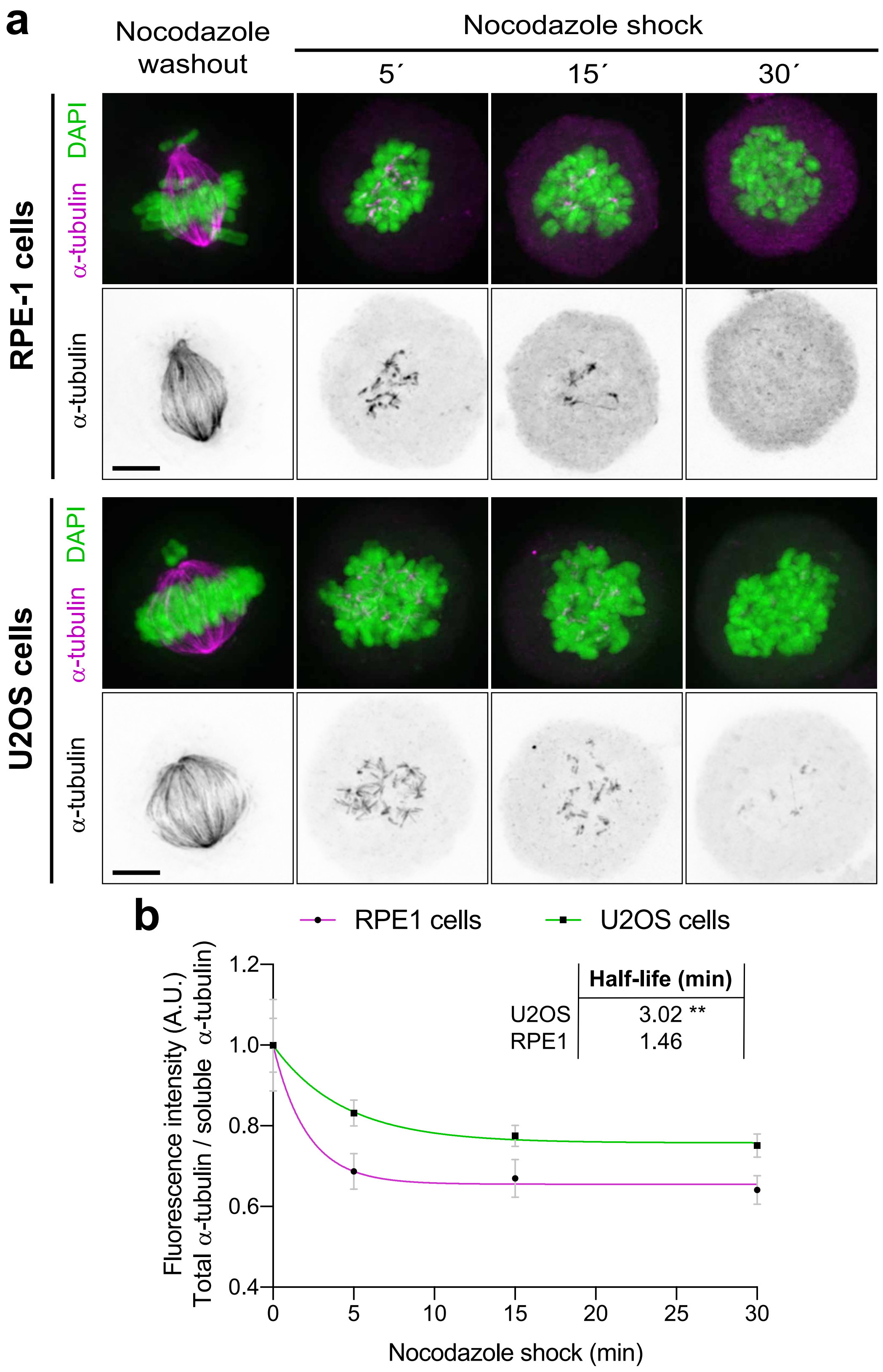


Gomes et al - Figure S7
