## Supplementary material for "Misaligned chromosomes that satisfy the spindle assembly checkpoint are a strong predictor of micronuclei formation in dividing cancer cells": Table S1

**Table S1.** Overview of the genes associated with chromosome congression defects analysed in the present study.

| Protein name | Subcelular localization | Chromosome congression defects described previously | Number of cells analyzed | Validation of depletion by WB | References |
|---|---|---|---|---|---|
| Nuf2 | Kinetochore | Prometaphase arrest; mitotic death | 310 cells from 2 independent experiments | yes | (DeLuca et al., 2005; DeLuca et al., 2002; Sundin et al., 2011) |
| Beclin-1 | kinetochore | Misaligned chromosomes; prometaphase arrest; mitotic death; | 374 cells from 2 independent experiments | no | (Fremont et al., 2013) |
| CLIP-170 | Kinetochore; mitotic spindle | Misaligned chromosomes; prometaphase arrest | 311 cells from 2 independent experiments | no | (Amin et al., 2015; Tanenbaum et al., 2006) |
| ATRX_oligo1 | Pericentromeric heterochromatin | Misaligned chromosomes; prometaphase arrest; abnormal nuclear morphology (lobulated nuclei and intranuclear DNA bridges); chromosome segregation defects (lagging chromosomes and chromosome bridges) | 392 cells from 2 independent experiments | yes | (Ritchie et al., 2008) |
| ATRX_oligo2 | Pericentromeric heterochromatin | Misaligned chromosomes; prometaphase arrest; abnormal nuclear morphology (lobulated nuclei and intranuclear DNA bridges); chromosome segregation defects (lagging chromosomes and chromosome bridges) | 400 cells from 2 independent experiments | yes | (Ritchie et al., 2008) |
| SPICE | Mitotic spindle; centrioles | Misaligned chromosomes; multipolar spindles | 504 cells from 2 independent experiments | no | (Archinti et al., 2010; Deretic et al., 2019) |
| CHICA | Mitotic spindle | Misaligned chromosomes, mitotic delay | 380 cells from 2 independent experiments | no | (Dunsch et al., 2012; Santamaria et al., 2008) |
| CDCA4 | | Multipolar spindle | 154 cells from 2 independent experiments | no | |
| Kif2a_oligo1 | Spindle poles | Partial knockdown of Kif2a accumulated cells with misaligned chromosomes; prometaphase delay; monopolar spindles | 600 cells from 3 independent experiments | yes | (Ganem and Compton, 2004; Jang et al., 2008) |
| Kif2a_oligo2 | Spindle poles | | 600 cells from 3 independent experiments | yes | (Ganem and Compton, 2004) |
| HIP1r | Mitotic spindle | Misaligned chromosomes; mitotic delay | 200 cells from 2 independent experiments | no | (Park, 2010) |
| Nucleophosmin (NPM1) | Chromosome periphery | Abnormal nuclear shape, tetraploid micronuclei formation; lost proliferation | 329 cells from 3 | no | (Amin et al., 2008) |

| | | | | | |
|---|---|---|---|---|---|
| | | ability; Misaligned chromosomes; disorganized spindles; mitotic delay | independent experiments | | |
| CENP-F/mitosin | kinetochore | Misaligned chromosomes; mitotic delay; aberrant spindle morphology, cell death | 398 cells from 2 independent experiments | yes | (Holt et al., 2005; Yang et al., 2005) |
| NudC | kinetochore | Misaligned chromosomes; prometaphase delay; Chromosome segregation defects (lagging chromosomes) | 338 cells from 2 independent experiments | no | (Chuang et al., 2013; Nishino et al., 2006) |
| RRS1 (Regulator of Ribosome Synthesis 1) | Chromosome periphery | Misaligned chromosomes; prometaphase delay | 397 cells from 2 independent experiments | no | (Gambe et al., 2009) |
| KIBRA | ND | Misaligned chromosomes; aberrant spindle morphology; Chromosome segregation defects (lagging chromosomes) | 401 cells from 2 independent experiments | no | (Zhang et al., 2012) |
| Nucleolin | Nucleoli; chromosome periphery | Misaligned chromosomes; prometaphase delay; aberrant spindle morphology | 150 cells from 2 independent experiments | no | (Li et al., 2009; Ma et al., 2007) |
| DDA3_oligo1 | Spindle microtubules; kinetochores; midbody | Misaligned chromosomes; prometaphase delay | 399 cells from 2 independent experiments | no | (Jang et al., 2010; Jang et al., 2011; Jang and Fang, 2011; Jang et al., 2008; Park et al., 2016) |
| DDA3_oligo2 | Spindle microtubules; kinetochores; midbody | | 383 cells from 2 independent experiments | no | (Jang et al., 2010; Jang et al., 2011; Jang and Fang, 2011; Jang et al., 2008; Park et al., 2016) |
| Bub1 | Kinetochore | Misaligned chromosomes; prometaphase delay; Chromosome segregation defects (lagging chromosomes) | 400 cells from 2 independent experiments | yes | (Johnson et al., 2004; Meraldi and Sorger, 2005; Morrow et al., 2005) |
| BubR1 | Kinetochore | Misaligned chromosomes; Chromosome segregation defects (lagging chromosomes) | 620 cells from 3 independent experiments | yes | (Ditchfield et al., 2003; Lampson and Kapoor, 2005) |
| Ska3_oligo1 | Kinetochore; mitotic spindle | Misaligned chromosomes, prometaphase delay, metaphase arrest, problems in maintenance of chromosome alignment, cohesion fatigue; cell death | 343 cells from 2 independent experiments | yes | (Raaijmakers et al., 2009) (Daum et al., 2009; Gaitanos et al., 2009; Sivakumar et al., 2014; Welburn et al., 2009) |
| Ska3_oligo2 | Kinetochore; mitotic spindle | | 400 cells from 2 independent experiments | yes | |
| TPX2 | Nucleus; spindle pole; spindle | Multipolar spindles; misaligned chromosomes | 403 cells from 3 independent experiments | no | (Garrett et al., 2002; Goshima, 2011) |
| Nup188_oligo1 | centrosomes | Misaligned chromosomes, prometaphase delay | 500 cells from 3 independent experiments | no | (Itoh et al., 2013) |
| Nup188_oligo2 | centrosomes | | 272 cells from 2 | no | |

| | | | | | |
|---|---|---|---|---|---|
| | | | independent experiments | | |
| Kif4a_oligo1 | Chromosome arms; spindle midzone | Misaligned chromosomes; mitotic delay; spindle defects; chromosome mis-segregation (lagging chromosomes and chromosome bridges) | 252 cells from 2 independent experiments | yes | (Mazumdar et al., 2004) |
| Kif4a_oligo2 | Chromosome arms; spindle midzone | | 348 cells from 2 independent experiments | yes | |
| m-calpain | ND | Misaligned chromosomes; prometaphase arrest; abnormal nuclear morphology, such as large and multiple lobed nuclei; Chromosome segregation errors (lagging chromosomes and chromosome bridges) | 200 cells from 1 experiment | no | (Honda et al., 2004; Magnaghi-Jaulin et al., 2010) |
| Zw10 | kinetochore | Misaligned chromosomes; prometaphase delay | 899 cells from 5 independent experiments | no | (Li et al., 2007b; Yang et al., 2007) |
| CENP-L | kinetochore | Chromosome alignment defects; chromosome mis-segregation (lagging chromosomes) | 405 cells from 3 independent experiments | no | (McHedlishvili et al., 2012) |
| NUSAP1 | Central spindle | Aberrant mitotic spindle, chromosome alignment defects; defective chromosome segregation; cytokinesis failure | 208 cells from 2 independent experiments | no | (Li et al., 2016; Raemaekers et al., 2003) |
| SAF-A/hnRNP-U | Spindle microtubules; spindle midzone | Aberrant mitotic spindles; Misaligned chromosomes; exit with misaligned chromosomes; prometaphase delay; cytokinesis failure | 375 cells from 2 independent experiments | no | (Ma et al., 2011) |
| Tastin/TROAP | Mitotic spindles in mitosis; centrosomes in interphase | Mitotic delay; Chromosome alignment defects; aberrant mitotic spindles; multipolar spindles | 304 cells from 2 independent experiments | no | (Yang et al., 2008) |
| Tankyrase-1 | Centrosome | Mitotic delay; metaphase delay | 566 cells from 3 independent experiments | no | (Chang et al., 2005; Dynek and Smith, 2004) |
| Ska1 | Kinetochore; mitotic spindle | Misaligned chromosomes, prometaphase delay, metaphase arrest, problems in maintenance of chromosome alignment, cohesion fatigue; cell death | 316 cells from 2 independent experiments | no | (Auckland et al., 2017; Gaitanos et al., 2009; Hanisch et al., 2006; Sivakumar et al., 2014; Welburn et al., 2009) |
| HURP | kinetochore | Misaligned chromosomes; prometaphase delay | 296 cells from 2 independent experiments | yes | (Sillje et al., 2006; Wong and Fang, 2006; Ye et al., 2011; Zhang et al., 2018) |
| Aki1/CC2D1A | centrosome | Multipolar spindles due to spindle pole fragmentation | 202 cells from 2 independent experiments | no | (Nakamura et al., 2009) |
| 4.1r | Mature centriole | Misaligned chromosomes; Multipolar spindles; Monopolar spindles | 451 cells from 3 independent experiments | yes | (Krauss et al., 2008) |
| HICE1/HAUS8 | Centrosome; mitotic spindle, spindle | Misaligned chromosomes; mitotic delay; aberrant mitotic | 377 cells from 2 | no | (Lawo et al., 2009; Wu et al., 2008) |

| | | | | | |
|---|---|---|---|---|---|
| | midzone; midbody | spindles; multipolar spindles, spindle pole fragmentation | independent experiments | | |
| Ska2 | Kinetochore; mitotic spindle | Misaligned chromosomes, prometaphase delay, metaphase arrest, problems in maintenance of chromosome alignment, cohesion fatigue | 399 cells from 2 independent experiments | yes | (Gaitanos et al., 2009; Hanisch et al., 2006; Sivakumar et al., 2014) |
| Kif18a | Plus-ends of kMTs | Misaligned chromosomes; prometaphase delay; long mitotic spindle; chromosomes mis-segregation (lagging chromosomes); Micronuclei formation | 330 cells from 2 independent experiments | no | (Fonseca et al., 2019; Huang et al., 2009; Liu et al., 2010; Mayr et al., 2007; Stumpff et al., 2008; Stumpff et al., 2012) |
| TACC3 | centrosome | Misaligned chromosomes; mitotic delay; disorganized spindles; cell death | 360 cells from 3 independent experiments | no | (Cheeseman et al., 2013; Gergely et al., 2003; Kimura et al., 2013; Lin et al., 2010; Schneider et al., 2007) |
| CLERC_Oligo1 | Centrosomes | Misaligned chromosomes; multipolar spindles | 294 cells from 2 independent experiments | no | (Muto et al., 2008) |
| CLERC_Oligo2 | Centrosomes | | 400 cells from 2 independent experiments | no | |
| Kizuna | Mature centriole; pericentriolar satellites | Misaligned chromosomes; multipolar spindles; spindle pole fragmentation | 358 cells from 2 independent experiments | no | (Oshimori et al., 2006) |
| ILK | Plasma membrane; focal adhesion; cytosol | Aberrant mitotic spindles, Misaligned chromosomes | 300 cells from 3 independent experiments | no | (Fielding et al., 2008) |
| Kinastrin/SKAP | Spindle pole; Kinetochore | Prometaphase delay; metaphase delay; multipolar spindles; spindle pole fragmentation | 355 cells from 2 independent experiments | no | (Dunsch et al., 2011; Fang et al., 2009; Huang et al., 2012; Schmidt et al., 2010) |
| Ninein | Mature centriole; pericentriolar satellites | Misaligned chromosomes; multipolar spindles | 200 cells from 2 independent experiments | no | (Logarinho et al., 2012) |
| NuMA | Nucleus; spindle pole | Aberrant mitotic spindles, Misaligned chromosomes | 188 cells from 2 independent experiments | yes | (Haren et al., 2009; Iwakiri et al., 2013) |
| Rae1_oligo1 | Nuclear pore; spindle pole | Misaligned chromosomes; multipolar spindles; lagging chromosomes; | 358 cells from 2 independent experiments | no | (Blower et al., 2005; Wong et al., 2006) |
| Rae1_oligo2 | | | 311 cells from 2 independent experiments | no | |
| RanBP2_oligo1 | Nuclear pore; kinetochore; spindle pole | Misaligned chromosomes; multipolar spindles | 400 cells from 2 independent experiments | no | (Joseph et al., 2004) |
| RanBP2_oligo2 | | | 231 cells from 1 experiment | no | |
| Spindly | Kinetochore; spindle pole | Misaligned chromosomes; prometaphase delay | 600 cells from 3 independent experiments | no | (Barisic et al., 2010; Raaijmakers et al., 2013) |

| | | | | | |
|---|---|---|---|---|---|
| STARD9 | Daughter centriole | Misaligned chromosomes; mitotic delay; multipolar spindles; spindle pole fragmentation; mitotic death | 198 cells from 2 independent experiments | no | (Torres et al., 2011) |
| CAMKIIγ (CAMK2G) | Cytosol | Misaligned chromosomes; multipolar spindles | 371 cells from 2 independent experiments | no | (Holmfeldt et al., 2005) |
| CENP-E | Kinetochore | Misaligned chromosomes; prometaphase delay; exit with misaligned chromosomes; die in or after mitosis | 259 cells from 3 independent experiments | yes | (Barisic et al., 2014; Maia et al., 2010; Stevens et al., 2011; Tanudji et al., 2004) |
| CHC17 clathrin | Mitotic spindle; centrosome | Chromosome misalignment; mitotic delay; disorganized spindles; multipolar spindles; spindle pole fragmentation | 299 cells from 2 independent experiments | no | (Foraker et al., 2012; Lin et al., 2010; Royle et al., 2005) |
| CEP72_oligo1 | centrosome | Misaligned chromosomes; multipolar spindles; spindle pole fragmentation | 379 cells from 2 independent experiments | yes | (Oshimori et al., 2009) |
| CEP72_oligo2 | centrosome | | 170 cells from 2 independent experiments | yes | |
| CEP90_oligo1 | Centrosome; pericentriolar satellites | Misaligned chromosomes; aberrant mitotic spindles; multipolar spindles | 471 cells from 3 independent experiments | yes | (Kim and Rhee, 2011) |
| CEP90_oligo2 | Centrosome; pericentriolar satellites | | 155 cells from 2 independent experiments | yes | |
| CLASP2b | Centrosome; kinetochore; microtubule plus ends; central spindle | Misaligned chromosomes; prometaphase delay; chromosome mis-segregation (lagging chromosomes and chromosome bridges); multipolar spindles; spindle pole fragmentation | 408 cells from 2 independent experiments | yes | (Girao et al., 2020; Logarinho et al., 2012; Mimori-Kiyosue et al., 2005) |
| CLASP1b | Centrosome; kinetochore; microtubule plus ends; central spindle | Misaligned chromosomes; chromosome mis-segregation (lagging chromosomes and chromosome bridges); multipolar spindles; spindle pole fragmentation | 402 cells from 2 independent experiments | yes | (Logarinho et al., 2012; Maiato et al., 2003; Mimori-Kiyosue et al., 2005) |
| CLASP1+2 | | | 300 cells from 2 independent experiments | yes | |
| Aurora B | Centromere; spindle, spindle midzone | Misaligned chromosomes; chromosome mis-segregation; cytokinesis defects | 287 cells from 2 independent experiments | yes | (Fuller et al., 2008; Hauf et al., 2003; Hegarat et al., 2011) |
| Astrin | Spindle pole; kinetochores | Misaligned chromosomes; prometaphase delay; multipolar spindles; spindle pole fragmentation | 400 cells from 2 independent experiments | yes | (Dunsch et al., 2011; Schmidt et al., 2010; Thein et al., 2007) |
| Aurora A | Centrosome; central spindle | Misaligned chromosomes; mitotic delay; multipolar spindles; spindle pole fragmentation; chromosome mis-segregation (lagging chromosomes and chromosome bridges) | 364 cells from 2 independent experiments | yes | (De Luca et al., 2008; De Luca et al., 2006; Hegarat et al., 2011; Hoar et al., 2007; Sasai et al., 2008) |

| | | | | | |
|---|---|---|---|---|---|
| INCENP | Centromere; spindle midzone | Prometaphase delay; Multipolar spindles; cytokinesis defects | 199 cells from 2 independent experiments | yes | (Mackay et al., 1998; Xu et al., 2009) |
| Haspin_oligo1 | Chromosome; centrosome | Misaligned chromosomes; mitotic arrest; prometaphase delay; multipolar spindles; spindle pole fragmentation | 399 cells from 2 independent experiments | no | (Dai et al., 2009; Dai et al., 2005) |
| Haspin_oligo2 | Chromosome; centrosome | | 401 cells from 2 independent experiments | no | |
| ARP1 | Kinetochore; mitotic spindle | Mitotic delay | 221 cells from 2 independent experiments | no | (Raaijmakers et al., 2013) |
| Spc25 | Kinetochore | Misaligned chromosomes; mitotic delay; aberrant mitotic spindles; multipolar spindles; cell death | 298 cells from 3 independent experiments | yes | (Bharadwaj et al., 2004; McCleland et al., 2004; Xu et al., 2014) |
| DYNLT3_oligo1 | kinetochore | Increased mitotic index, particularly the number of cells in prophase/prometaphase | 309 cells from 2 independent experiments | no | (Lo et al., 2007) |
| DYNLT3_oligo2 | | | 564 cells from 3 independent experiments | no | |
| DYNLRB1/Road block-1 | Kinetochore; mitotic spindle | Misaligned chromosomes; prometaphase delay | 165 cells from 2 independent experiments | no | (Raaijmakers et al., 2013) |
| Spc24 | Kinetochore | Misaligned chromosomes | 399 cells from 2 independent experiments | yes | (Bharadwaj et al., 2004; Xu et al., 2014) |
| NdeL1 | Kinetochore; mitotic spindle | Misaligned chromosomes; prometaphase delay; chromosome mis-segregation | 462 cells from 2 independent experiments | no | (Raaijmakers et al., 2013; Vergnolle and Taylor, 2007) |
| LIS1/PAFAH1B1 | Kinetochore; mitotic spindle | Misaligned chromosomes; prometaphase delay; chromosome mis-segregation (lagging chromosomes and chromosome bridges) | 469 cells from 3 independent experiments | no | (Moon et al., 2014; Raaijmakers et al., 2013) |
| HSET/KIF1C | Microtubules | | 400 cells from 2 independent experiments | no | (Auckland and McAinsh, 2015) |
| Nde1 | Kinetochore; mitotic spindle | Misaligned chromosomes; prometaphase delay; | 611 cells from 3 independent experiments | yes | (Raaijmakers et al., 2013; Vergnolle and Taylor, 2007) |
| DLIC2 | Spindle pole | Misaligned chromosomes; prometaphase delay | 400 cells from 2 independent experiments | no | (Horgan et al., 2011; Raaijmakers et al., 2013) |
| HAUS6_oligo1 | Centrosome, spindle | Misaligned chromosomes; aberrant mitotic spindles; spindle pole fragmentation; mitotic delay | 345 cells from 2 independent experiments | yes | (Lawo et al., 2009) |
| HAUS6_oligo2 | Centrosome, spindle | | 168 cells from 1 experiment | yes | |
| HAUS1_oligo1 | Centrosome, spindle | Misaligned chromosomes; aberrant mitotic spindles; | 400 cells from 2 | no | (Einarson et al., 2004; Lawo et al., 2009) |

| | | | | | |
|---|---|---|---|---|---|
| | | spindle pole fragmentation; mitotic delay | independent experiments | | |
| HAUS1_oligo2 | Centrosome, spindle | | 396 cells from 2 independent experiments | no | |
| DIC2/DYNC1I2 | Kinetochore; mitotic spindle | Misaligned chromosomes; prometaphase delay | 184 cells from 2 independent experiments | no | (Raaijmakers et al., 2013) |
| survivin | Centromeres; spindle midzone | Misaligned chromosomes; prometaphase delay; chromosome segregation errors; Cytokinesis failure, mitotic catastrophe | 294 cells from 2 independent experiments | yes | (Carvalho et al., 2003; Lens et al., 2003; Uren et al., 2000) |
| CPAP/ CENP-J | centriole | Misaligned chromosomes; mitotic delay; multipolar spindles; apoptosis | 215 cells from 2 independent experiments | no | (Cho et al., 2006) |
| Hec1 | kinetochore | Misaligned chromosomes; mitotic arrest | 191 cells from 2 independent experiments | yes | (Joseph et al., 2004; Lee et al., 2011; Li et al., 2007a; Martin-Lluesma et al., 2002; Sundin et al., 2011) |
| Borealin | Centromere; spindle midzone | | 368 cells from 2 independent experiments | no | (Gassmann et al., 2004) |
| Scc1/Rad21 | Chromosome; centrosome | Misaligned chromosomes; prometaphase delay; multipolar spindles; spindle pole fragmentation | 384 cells from 2 independent experiments | no | (Beauchene et al., 2010; Dai et al., 2009; Diaz-Martinez et al., 2010) |
| Myosin 10 | Spindle pole | Misaligned chromosomes; mitotic delay; aberrant mitotic spindles; spindle pole fragmentation; cytokinesis failure | 500 cells from 3 independent experiments | no | (Woolner et al., 2008) |
| Sgo1/Shugoshin | Centromere; Kinetochore; centrosome; spindle pole | Misaligned chromosomes; prometaphase delay; spindle pole fragmentation | 414 cells from 2 independent experiments | no | (McGuinness et al., 2005; Wang et al., 2008) |
| HAUS3_oligo1 | Centrosome, spindle | Misaligned chromosomes; aberrant mitotic spindles; spindle pole fragmentation; | 400 cells from 2 independent experiments | no | (Lawo et al., 2009) |
| HAUS3_oligo2 | Centrosome, spindle | | 400 cells from 2 independent experiments | no | |
| DHC/DYNC1H1 | Kinetochore; mitotic spindle | Misaligned chromosomes; prometaphase delay; aberrant mitotic spindles | 396 cells from 2 independent experiments | yes | (Barisic et al., 2014; Raaijmakers et al., 2013) |
| CENP-T | kinetochore | Misaligned chromosomes, prometaphase arrest; multipolar spindles; cell death | 395 cells from 2 independent experiments | no | (McKinley et al., 2015; Prendergast et al., 2011; Wood et al., 2016) |
| MLL | Mitotic spindle | Misaligned chromosomes; mitotic delay | 560 cells from 3 independent experiments | no | (Ali et al., 2017) |
| CENP-W | kinetochore | Misaligned chromosomes; mitotic delay; multipolar spindles | 380 cells from 2 independent experiments | no | (Chun et al., 2016; Prendergast et al., 2011) |

| | | | | | |
|---|---|---|---|---|---|
| Shp2_oligo1 | Kinetochore; centrosome; spindle midzone; midbody | Misaligned chromosomes; mitotic delay; chromosome mis-segregation (lagging chromosomes) | 528 cells from 3 independent experiments | yes | (Liu et al., 2012) |
| Shp2_oligo2 | | | 455 cells from 3 independent experiments | yes | |
| ASURA/PHB2 | cytoplasm | Misaligned chromosomes; mitotic arrest | 400 cells from 2 independent experiments | no | (Equilibrina et al., 2013; Lee et al., 2011; Takata et al., 2007) |
| CENP-H | kinetochore | Misaligned chromosomes; aberrant mitotic spindles; multipolar spindles | 324 cells from 3 independent experiments | no | (Amaro et al., 2010; Orthaus et al., 2006) |
| Kif14_oligo1 | Spindle poles; mitotic spindle; midbody | Misaligned chromosomes; cytokinesis failure; binucleated cells | 436 cells from 3 independent experiments | no | (Carleton et al., 2006; Zhu et al., 2005) |
| Kif14_oligo2 | Spindle poles; mitotic spindle; midbody | | 277 cells from 2 independent experiments | no | |
| WDR5 | Mitotic spindle | Misaligned chromosomes; mitotic delay | 383 cells from 2 independent experiments | no | (Ali et al., 2017) |
| TAO1/MARKK | microtubules | Misaligned chromosomes; prometaphase delay; Muti-lobed nuclei; chromosome mis-segregation (lagging chromosomes) | 416 cells from 2 independent experiments | no | (Draviam et al., 2007; Shrestha et al., 2014) |
| Nup88 | Mitotic spindle | Misaligned chromosomes; multipolar spindles | 300 cells from 2 independent experiments | no | (Hashizume et al., 2010) |
| ASB7 | ND | Misaligned chromosomes | 300 cells from 2 independent experiments | no | (Uematsu et al., 2016) |
| And-1_oligo1 | cytoplasm | Misaligned chromosomes; prometaphase delay | 415 cells from 2 independent experiments | yes | (Jaramillo-Lambert et al., 2013) |
| And-1_oligo2 | cytoplasm | | 168 cells from 1 experiment | yes | |
| Septin-7 | Spindle poles; mitotic spindle; midbody | Misaligned chromosomes; mitotic arrest | 222 cells from 2 independent experiments | no | (Zhu et al., 2008) |
| ANKRD53 | Spindle poles | Misaligned chromosomes; mitotic delay; Multinucleated cells | 360 cells from 2 independent experiments | no | (Kim and Jang, 2016) |
| TRAMM | Perinuclear region | Misaligned chromosomes; mitotic delay | 371 cells from 2 independent experiments | no | (Milev et al., 2015) |
| Septin-2_oligo1 | plasma membrane, cleavage furrow and midbody | Misaligned chromosomes; mitotic delay; binucleated cells | 411 cells from 2 independent experiments | yes | (Spiliotis et al., 2005) |

| | | | | | |
|---|---|---|---|---|---|
| Septin-2_oligo2 | | | 325 cells from 2 independent experiments | yes | |
| Seh1 | kinetochores | Misaligned chromosomes; mitotic delay; cytokinesis defects | 200 cells from 2 independent experiments | no | (Platani et al., 2009; Zuccolo et al., 2007) |
| CENP-Q_oligo1 | kinetochore | Misaligned chromosomes; prometaphase delay | 430 cells from 3 independent experiments | no | (Bancroft et al., 2015) |
| CENP-Q_oligo2 | kinetochore | | 352 cells from 3 independent experiments | no | |
| NF-1 | Astral microtubules; mitotic spindle, centrosomes; midbody | Misaligned chromosomes | 375 cells from 2 independent experiments | no | (Koliou et al., 2016) |
| Scramble RNAi | | | 7229 cells from 45 independent experiments | | |
| Nup107 | kinetochore | Misaligned chromosomes; mitotic delay; cytokinesis defects | 428 cells from 3 independent experiments | no | (Platani et al., 2009; Zuccolo et al., 2007) |
| Usp16 | Cytoplasmic in interphase; kinetochore | Misaligned chromosomes; prometaphase delay | 663 cells from 3 independent experiments | no | (Zhuo et al., 2015) |
| NDR1 | ND | Misaligned chromosomes | 264 cells from 2 independent experiments | no | (Oh et al., 2010) |
| GAK_oligo1 | Trans-Golgi network | Misaligned chromosomes; prometaphase arrest; multipolar spindles; spindle pole fragmentation | 333 cells from 2 independent experiments | yes | (Shimizu et al., 2009) |
| GAK_oligo2 | Trans-Golgi network | | 376 cells from 3 independent experiments | yes | |
| HAUS7 | Centrosome, spindle | Misaligned chromosomes; aberrant mitotic spindles; spindle pole fragmentation; | 303 cells from 2 independent experiments | no | (Lawo et al., 2009) |
| CENP-M | kinetochore | Misaligned chromosomes | 599 cells from 3 independent experiments | no | (Basilico et al., 2014; Foltz et al., 2006) |
| CENP-U | kinetochore | Misaligned chromosomes; chromosome mis-segregation (lagging chromosomes) | 278 cells from 3 independent experiments | no | (Hua et al., 2011) |
| MST1 | ND | Misaligned chromosomes; prometaphase delay; cell death | 300 cells from 2 independent experiments | no | (Oh et al., 2010) |
| PTEN | Centrosome; mitotic spindle; midbody | Misaligned chromosomes; mitotic delay; spindle pole fragmentation; mitotic catrastrophe | 280 cells from 2 independent experiments | no | (He et al., 2016) |

| | | | | | |
|---|---|---|---|---|---|
| HAUS2 | Centrosome, spindle | Misaligned chromosomes; aberrant mitotic spindles; spindle pole fragmentation; | 339 cells from 3 independent experiments | no | (Lawo et al., 2009) |
| CENP-N | kinetochore | Misaligned chromosomes, multipolar spindles | 300 cells from 2 independent experiments | no | (McKinley et al., 2015) |
| Hsp72 | Mitotic spindle; midbody | Misaligned chromosomes; prometaphase delay | 428 cells from 3 independent experiments | no | (O'Regan et al., 2015) |
| HAUS5 | Centrosome, spindle | Misaligned chromosomes; aberrant mitotic spindles; spindle pole fragmentation; | 397 cells from 2 independent experiments | no | (Lawo et al., 2009) |
| HAUS4_oligo1 | Centrosome, spindle | Misaligned chromosomes; aberrant mitotic spindles; spindle pole fragmentation; | 324 cells from 2 independent experiments | no | (Lawo et al., 2009) |
| HAUS4_oligo2 | | | 254 cells from 2 independent experiments | no | |
| DYNLT1 | Kinetochore; mitotic spindle | ND | 596 cells from 3 independent experiments | no | |
| Rab5 | Early endosomes | Misaligned chromosomes; prometaphase delay | 579 cells from 3 independent experiments | no | (Serio et al., 2011) |
| CENP-I | Kinetochore | Misaligned chromosomes, multipolar spindles; chromosome mis-segregation; apoptosis | 305 cells from 2 independent experiments | yes | (Liu et al., 2003; McKinley et al., 2015; Nishihashi et al., 2002) |
| Dsn1 | Kinetochore | Misaligned chromosomes; prometaphase delay | 582 cells from 3 independent experiments | yes | (Kline et al., 2006) |
| CENP-P | kinetochore | Misaligned chromosomes; mitotic delay | 567 cells from 3 independent experiments | no | (Bancroft et al., 2015) |
| ZwinT | Kinetochore | Misaligned chromosomes; cell death; chromosome mis-segregation (lagging chromosomes and chromosome bridges) | 410 cells from 2 independent experiments | no | (Lin et al., 2006; Wang et al., 2004) |
| Nsl1 | Kinetochore | Misaligned chromosomes | 300 cells from 2 independent experiments | no | (Kline et al., 2006) |
| Mis12 | Kinetochore | Misaligned chromosomes | 200 cells | no | (Kline et al., 2006) |
| KNL1 | Kinetochore | Misaligned chromosomes | 291 cells from 2 independent experiments | no | (Caldas and DeLuca, 2014; Ghongane et al., 2014) |
| Nup153 | ND | Multilobed nuclei; cytokinesis abnormalities | 200 cells from 2 independent experiments | no | (Chatel and Fahrenkrog, 2011; Lussi et al., 2010; Mackay et al., 2009) |
