## Supplementary material for "Misaligned chromosomes that satisfy the spindle assembly checkpoint are a strong predictor of micronuclei formation in dividing cancer cells": Table S2

**Table S2.** List all reagents used, respective sources and siRNA depletion conditions.

| Reagent type | Designation | Source or reference | Identifiers | Additional information |
|---|---|---|---|---|
| Cell line (human) | HeLa H2b-GFP, α-tubulin-mRFP | Generated by lentiviral transduction | | |
| Cell line (human) | HeLa parental | Provided by Y.Mimori-Kiyosie | | |
| Cell line (human) | HeLa Mad2-GFP | Schweizer et al., 2013 | | |
| Cell line (human) | Cyclin B1-Venus HeLa | Provided by J.Pines | | |
| Cell line (human) | U2OS parental | Provided by S. Geley | | |
| Cell line (human) | U2OS H2B-GFP, α-tubulin-cherry | Provided by S. Geley | | |
| Cell line (human) | hTERT-REP1 (RPE1) parental | Provided by Ben Black | | |
| Cell line (human) | RPE1 H2B-GFP, α-tubulin-mCherry | Generated by lentiviral transduction | | |
| Antibody | mouse anti-Mad1 | Merck Millipore | Cat#MABE867 | IF (1:500) |
| Antibody | mouse anti-α-tubulin | Sigma | T5168 | IF (1:2000)<br>WB (1:5000) |
| Antibody | rabbit anti-β-tubulin | Abcam | Ab6046 | IF (1:2000) |
| Antibody | guinea pig anti-CENP-C | MBL International | PD030 | IF (1:1000) |
| Antibody | mouse anti-Aim1 | BD Biosciences | Cat#611083 | WB (1:1000) |
| Antibody | mouse anti-Hec1 (9GA) | Abcam | Ab3613 | WB (1:1000) |
| Antibody | mouse anti-Dsn1 | Provided by A. Musacchio | | WB (1:500) |
| Antibody | mouse anti-ATRX | Santa Cruz Biotechnology | sc-55584 | WB (1:1000) |
| Antibody | mouse anti-CENP-F | BD Biosciences | 610768 | WB (1:1000) |
| Antibody | rabbit anti-CEP72 | Novus Biologicals | NB100-60661 | WB (1:1000) |
| Antibody | mouse anti-GAK | R&D Systems | MAB6918 | WB (1:500) |
| Antibody | rabbit anti-WDHD1/And-1 | Novus Biologicals | NBP1-89091 | WB (1:1000) |
| Antibody | rabbit anti-Aurora A | Novus Biologicals | NB100-267 | WB (1:1000) |
| Antibody | rabbit anti-HURP | a gift from P. Meraldi | | WB (1:200) |
| Antibody | mouse anti-INCENP | Santa Cruz Biotechnology | sc-376514 | WB (1:500) |
| Antibody | rabbit anti-LRRCC1/CLERC | Abcam | ab95450 | WB (1:1000) |
| Antibody | mouse anti-Sgo1 (F-8) | Santa Cruz Biotechnology | sc-393993 | WB (1:1000) |
| Antibody | rabbit anti-DHC | ThermoFisher Scientific | PA5-49373 | WB (1:500) |
| Antibody | mouse anti-Nde1 | Abnova | H00054820-M01 | WB (1:500) |
| Antibody | sheep anti-Bub1 | a gift from Stephen Taylor | | WB (1:1000) |
| Antibody | rabbit anti-Septin-2 | Novus Biologicals | NBP1-85212 | WB (1:500) |
| Antibody | mouse anti-Ska3 | Santa Cruz Biotechnology | sc-390326 | WB (1:1000) |
| Antibody | rabbit anti-CEP90 | Novus Biologicals | NBP2-56805 | WB (1:1000) |
| Antibody | mouse anti-ska2 | Santa Cruz Biotechnology | sc-514495 | WB (1:1000) |
| Antibody | mouse anti-4.1r (B-11) | Santa Cruz Biotechnology | sc-166759 | WB (1:1000) |
| Antibody | rabbit anti-Astrin (N-terminal) | a gift from Duane Compton | | WB (1:1000) |
| Antibody | rabbit anti-Kif4a | ThermoFisher Scientific | pa5-30492 | WB (1:1000) |
| Antibody | rat anti-CLASP1 | Maffini et al 2009 | | WB (1:50) |

| | | | | |
|---|---|---|---|---|
| Antibody | rat anti-CLASP2 | Maffini et al 2009 | | WB (1:50) |
| Antibody | rabbit anti-BubR1 | Abcam | ab 200062 | WB (1:500) |
| Antibody | mouse anti-Nde1 | Abnova | H00054820-M01 | WB (1:1000) |
| Antibody | rabbit anti-SHP2 | Abcam | ab10555 | WB (1:1000) |
| Antibody | rabbit anti-survivin | Novus Biologicals | NB500-201 | WB (1:1000) |
| Antibody | mouse anti-GAPDH | Proteintech | 60004-1-Ig | WB (1:40000) |
| Antibody | rabbit anti-vinculin | ThermoFisher Scientific | 700062 | WB (1:1000) |
| Antibody | anti-mouse-HRP | Jackson ImmunoResearch Laboratories | | WB (1:1500) |
| Antibody | anti-rabbit-HRP | Jackson ImmunoResearch Laboratories | | WB (1:1500) |
| Antibody | anti-sheep-HRP | Jackson ImmunoResearch Laboratories | | WB (1:1500) |
| Antibody | anti-rat-HRP | Jackson ImmunoResearch Laboratories | | WB (1:1500) |
| Chemical compound, drug | nocodazole | Sigma-Aldrich | Cat#M1401 | 1 µM |
| Chemical compound, drug | MG132 | EMD Millipore | Cat#133407-82-6 | 5 µM |
| Oligonucleotide | 5´- CUUCCUCUCUUUCUCUCCCUUGUGATT-3´ (scramble) | Sigma-Aldrich | | |
| Oligonucleotide | 5´-CCAGGACUACGAGGCGCUGTT-3´ (m-calpain) | Sigma-Aldrich | (Honda et al., 2004) | 50nM, 96h |
| Oligonucleotide | 5´-AAGCAUGCCGUGAAACGUAUATT-3´ (Nuf2) | Sigma-Aldrich | (DeLuca et al., 2002) | 50nM, 24h |
| Oligonucleotide | 5´-GCUCAGUAUCAGAGAGAAUTT-3´ (Beclin-1) | Sigma-Aldrich | (Fremont et al., 2013) | 50nM, 48h |
| Oligonucleotide | 5´-GCACAGCUCUGAAGACACCTT-3´ (CLIP-170) | Sigma-Aldrich | (Tanenbaum et al., 2006) | 50nM, 96h |
| Oligonucleotide | 5´-GAGGAAACCUUCAAUUGUATT-3´ (ATRX_Oligo1) | Sigma-Aldrich | (Ritchie et al., 2008) | 50nM, 72h |
| Oligonucleotide | 5´-GCAGAGAAAUUCCUAAAGATT-3´ (ATRX_Oligo2) | Sigma-Aldrich | (Ritchie et al., 2008) | 50nM, 72h |
| Oligonucleotide | 5´-GCUGAGAACAAAUGAGUCATT-3´ (SPICE) | Sigma-Aldrich | (Archinti et al., 2010) | 50nM, 48h |
| Oligonucleotide | 5´-CCAGGAUAGCAAGCUCUCAAATT-3´ (CHICA) | Sigma-Aldrich | (Santamaria et al., 2008) | 50nM, 48h |
| Oligonucleotide | 5´-GCUGCAUGGAAGAGCUGUUTT-3´ (CDCA4) | Sigma-Aldrich | (Wang et al., 2008) | 50nM, 96h |
| Oligonucleotide | 5´-GGAAUGGCAUCCUGUGAAATT-3´ (Kif2a_Oligo1) | Sigma-Aldrich | (Jang et al., 2008) | 50nM, 72h |
| Oligonucleotide | 5´-GGCAAAGAGAUUGACCUGGTT-3´ (Kif2a_Oligo 2) | Sigma-Aldrich | (Ganem and Compton, 2004) | 50nM, 72h |
| Oligonucleotide | 5´-UUCUCAUGAUGCGUGCCCAGGAUGATT-3´ (HIP1r) | Sigma-Aldrich | (Park, 2010) | 50nM, 96h |
| Oligonucleotide | 5´-AGAUGAUGAUGAUGAUGAUUUTT-3´ (Nucleophosmin) | Sigma-Aldrich | (Amin et al., 2008) | 50nM, 48h |
| Oligonucleotide | 5´-AAGAGAAGACCCCAAGUCAUCTT-3´ (CENP-F) | Sigma-Aldrich | (Holt et al., 2005) | 50nM, 72h |
| Oligonucleotide | 5´-AACAGACUUUUUCAUUGGAGGTT-3´ (NudC) | Sigma-Aldrich | (Nishino et al., 2006) | 50nM, 48h |
| Oligonucleotide | 5´-CUACCGGACACCAGAGUAATT-3´ (RRS1) | Sigma-Aldrich | (Gambe et al., 2009) | 50nM, 72h |
| Oligonucleotide | 5´-GGUUGGAGAUUACUUCAUATT-3´ (KIBRA) | Sigma-Aldrich | (Zhang et al., 2012) | 50nM,96h |
| Oligonucleotide | 5´-AGAGUUUGCUUCAUUCGAATT-3´ (Nucleolin) | Sigma-Aldrich | | 50nM, 96h |
| Oligonucleotide | 5´-AAGCAAGACUUCAGUAGCATT-3´ (DDA3_Oligo1) | Sigma-Aldrich | (Jang et al., 2008) | 50nM, 72h |
| Oligonucleotide | 5´-CCACCGAAGTGACCCAAATTT-3´ (DDA3_Oligo 2) | Sigma-Aldrich | (Jang et al., 2008) | 50nM, 24h |
| Oligonucleotide | 5´-AAAUACCACAAUGACCCAAGATT-3´ (Bub1) | Sigma-Aldrich | (Johnson et al., 2004) | 50nM, 48h |
| Oligonucleotide | 5´-AACGGGCAUUUGAAUAUGAAATT-3´ (BubR1) | Sigma-Aldrich | (Ditchfield et al., 2003) | 50nM, 48h |

| | | | | |
|---|---|---|---|---|
| Oligonucleotide | 5´-AAUCCAGGCUCAAUGAUAATT-3´ (Ska3_Oligo1) | Sigma-Aldrich | (Raaijmakers et al., 2009) | 50nM, 24h |
| Oligonucleotide | 5´-AGACAAACAUGAACAUUAATT-3´ (Ska3_Oligo2) | Sigma-Aldrich | (Gaitanos et al., 2009) | 50nM, 48h |
| Oligonucleotide | 5´-AAGGAGAUACUCAAAACAUAGTT-3´ (TPX2) | Sigma-Aldrich | (Garrett et al., 2002) | 50nM, 96h |
| Oligonucleotide | 5´-AUUUCUAGCAGCAUGGACUGUUCCCTT-3´ (Nup188_Oligo1) | Sigma-Aldrich | (Itoh et al., 2013) | 50nM, 96h |
| Oligonucleotide | 5´-GGUAGUAGGCAGACCAAUATT-3´ (Nup188_Oligo2) | Sigma-Aldrich | (Labade et al., 2016) | 50nM, 96h |
| Oligonucleotide | 5´-GCAAUUGAUUACCCAGUUATT-3´ (Kif4a_Oligo1) | Sigma-Aldrich | (Mazumdar et al., 2004) | 50nM, 96h |
| Oligonucleotide | 5´-GAAAGATCCTGGCTCAAGATT-3´ (Kif4a_Oligo2) | Sigma-Aldrich | (Mazumdar et al., 2004) | |
| Oligonucleotide | 5´-UGAUCAAUGUGCUGUUCAATT-3´ (Zw10) | Sigma-Aldrich | (Kops et al., 2005) | 50nM, 72h |
| Oligonucleotide | 5´-CCAUUAUGUGGCUACUACUGAAUUU-3´ (CENP-L) | Sigma-Aldrich | (McHedlishvili et al., 2012) | 50nM, 96h |
| Oligonucleotide | 5´-AAGCACCAAGAAGCUGAGAAUTT-3´ (NUSAP1) | Sigma-Aldrich | (Raemaekers et al., 2003) | 50nM, 96h |
| Oligonucleotide | 5´-GAACUCUCGUAUGCUAAGATT-3´ (SAF-A) | Sigma-Aldrich | (Ma et al., 2011) | 50nM, 24h |
| Oligonucleotide | 5´-GCCUGAUCUUCUCUUCCCATT-3´ (Tastin) | Sigma-Aldrich | (Yang et al., 2008) | 50nM, 96h |
| Oligonucleotide | 5´-AACAAUUCACCGUCGUCCUCUTT-3´ (Tankyrase-1) | Sigma-Aldrich | (Dynek and Smith, 2004) | 50nM, 72h |
| Oligonucleotide | 5´-CCCGCUUAACCUAUAAUCAAATT-3’ (Ska1) | Sigma-Aldrich | (Hanisch et al., 2006) | 50nM, 48h |
| Oligonucleotide | 5´-AAUGACUCGAUCAGCUACUCATT-3´ (HURP) | Sigma-Aldrich | (Sillje et al., 2006) | 50nM, 48h |
| Oligonucleotide | 5´-CCCUGGCGAUCUGGAUGUCUUUGUU-3´ (Aki) | Sigma-Aldrich | (Nakamura et al., 2009) | 50nM, 96h |
| Oligonucleotide | 5´-GAAAGUCUGUGUAGAACAUUU-3´ (4.1r) | Sigma-Aldrich | (Krauss et al., 2008) | 50nM, 72h |
| Oligonucleotide | 5´-GGGAGAACUUGAUGUUGGUGAUUCGTT-3´ (HICE1) | Sigma-Aldrich | (Wu et al., 2008) | 50nM, 48h |
| Oligonucleotide | 5´-AAGAAAUCAAGACUAAUCAUCTT-3´ (Ska2) | Sigma-Aldrich | (Hanisch et al., 2006) | 50nM, 72h |
| Oligonucleotide | 5´-ACCAACAACAGUGCCAUAAATT-3´ (Kif18a) | Sigma-Aldrich | (Huang et al., 2009) | 50nM, 24h |
| Oligonucleotide | 5´-CACGGGCGCGGAGGUGGAUUATT-3´ (TACC3) | Sigma-Aldrich | (Fielding et al., 2011) | 50nM, 48h |
| Oligonucleotide | 5´-GGAGAAAGAUGGAGACGAUTT-3´ (CLERC_Oligo1) | Sigma-Aldrich | (Muto et al., 2008) | 50nM, 24h |
| Oligonucleotide | 5´-CAGAUAGGCUAAAGGAAAUTT-3´ (CLERC_Oligo2) | Sigma-Aldrich | (Muto et al., 2008) | 50nM, 72h |
| Oligonucleotide | 5´-AAGCGAUUUGAGCGUGUCCAATT-3´ (Kizuna) | Sigma-Aldrich | (Oshimori et al., 2006) | 50nM, 72h |
| Oligonucleotide | 5´-AAGACGCUCAGCAGACAUGUGGATT-3´ (ILK) | Sigma-Aldrich | (Fielding et al., 2011) | 50nM, 96h |
| Oligonucleotide | 5´-AGGCUACAAACCACUGAGUAATT-3´ (Kinastrin) | Sigma-Aldrich | (Dunsch et al., 2011) | 50nM, 96h |
| Oligonucleotide | 5´-UAUGAGCAUUGAGGCAGAGTT-3´ (Ninein) | Sigma-Aldrich | (Logarinho et al., 2012) | 50nM, 96h |
| Oligonucleotide | 5´-GGCGUGGCAGGAGAAGUUCTT-3´ (NuMA) | Sigma-Aldrich | (Logarinho et al., 2012) | 50nM, 96h |
| Oligonucleotide | 5´-GCAGUAACCAAGCGAUACATT-3´ (Rae1_Oligo1) | Sigma-Aldrich | (Wong et al., 2006) | 50nM, 72h |
| Oligonucleotide | 5´-GAGUUGCUAUUCACUAUAUTT-3´ (Rae1_Oligo2) | Sigma-Aldrich | (Blower et al., 2005) | 50nM, 48h |
| Oligonucleotide | 5´-AAGGACAGUGGGAUUGUAGUGTT-3´(RanBP2_Oligo1) | Sigma-Aldrich | | 50nM, 72h |
| Oligonucleotide | 5´-AACAACACCAAAAGCAGUGGUTT-3´ (RanBP2_Oligo2) | Sigma-Aldrich | | 50nM, 72h |
| Oligonucleotide | 5´-GAAAGGGUCUCAAACUGAATT-3´ (Spindly) | Sigma-Aldrich | (Barisic et al., 2010) | 50nM, 48h |
| Oligonucleotide | 5´-GAGUUGCCAAAGGCUAUAATT-3´ (STARD9) | Sigma-Aldrich | (Srivastava and Panda, 2018) | 50nM, 72h |
| Oligonucleotide | 5´-GCAGAUGCCAGCCACUGUATT-3´ (CAMKIIγ) | Sigma-Aldrich | SASI_Hs01_00118118 | 50nM, 24h |
| Oligonucleotide | 5´-GAACUAAGAAGAAGCGUAUTT-3´ (CENP-E) | Sigma-Aldrich | (Maia et al., 2010) | 50nM, 24h |

| | | | | |
|---|---|---|---|---|
| Oligonucleotide | 5´-AAGCAAUGAGCUGUUUGAAGATT-3´ (CHC17 clatrin) | Sigma-Aldrich | (Vassilopoulos et al., 2009) | 50nM, 96h |
| Oligonucleotide | 5´-UUGCAGAUCGCUGGACUUCAATT-3´ (CEP72_Oligo1) | Sigma-Aldrich | (Oshimori et al., 2009) | 50nM, 72h |
| Oligonucleotide | 5´-GAGUUUAACAGGUCUGAAATT-3´ (CEP72_Oligo2) | Sigma-Aldrich | SASI_Hs02_00351368 | 50nM, 96h |
| Oligonucleotide | 5´-GCAGCUGACAGAGACAUAUTT-3´ (CEP90_Oligo1) | Sigma-Aldrich | (Kim and Rhee, 2011) | 50nM, 48h |
| Oligonucleotide | 5´-CACCUUAGAGCAAACUGUUTT-3´ (CEP90_Oligo2) | Sigma-Aldrich | SASI_HS01_00230208 | 50nM, 96h |
| Oligonucleotide | 5´-GGAUGAUUUACAAGACUGGTT-3´ (CLASP1b) | Sigma-Aldrich | (Mimori-Kiyosue et al., 2005) | 50nM, 48h |
| Oligonucleotide | 5´-GACAUACAUGGGUCUUAGATT-3´ (CLASP2b) | Sigma-Aldrich | (Mimori-Kiyosue et al., 2005) | 50nM, 48h |
| Oligonucleotide | 5´-AACGCGGCACUUCACAAUUGATT-3´ (Aurora B) | Sigma-Aldrich | (Fuller et al., 2008) | 50nM, 24h |
| Oligonucleotide | 5´-UCCCGACAACUCACAGAGAAATT-3´ (Astrin) | Sigma-Aldrich | (Thein et al., 2007) | 50nM, 48h |
| Oligonucleotide | 5´-AUGCCCUGUCUUACUGUCATT-3´ (Aurora A) | Sigma-Aldrich | (Kuang et al., 2017) | 50nM, 24h |
| Oligonucleotide | 5´-CUCAGAAGAACCGACGGAATT-3´ (INCENP) | Sigma-Aldrich | SASI_Hs01_00219348 | 50nM, 24h |
| Oligonucleotide | 5´-GGCUUUAUCGGGCUGAACUTT-3´ (Haspin_Oligo1) | Sigma-Aldrich | SASI_Hs02_00359157 | 50nM, 48h |
| Oligonucleotide | 5´- GCUUUGAGCACCGAGACUUTT-3´ (Haspin_Oligo2) | Sigma-Aldrich | SASI_Hs01_00243245 | 50nM, 72h |
| Oligonucleotide | 5´- CCUUCAAUGUGCCCGCUCUTT-3´ (ARP1) | Sigma-Aldrich | SASI_Hs01_00015229 | 50nM, 48h |
| Oligonucleotide | 5´- CUGCAAAUAUCCAGGAUCUTT-3´ (Spc25) | Sigma-Aldrich | SASI_Hs01_00193697 | 50nM, 24h |
| Oligonucleotide | 5´- CAUAGUAAUUGGCAGAUUATT-3´ (DYNLT3_Oligo1) | Sigma-Aldrich | SASI_Hs01_00147343 | 50nM, 96h |
| Oligonucleotide | 5´- GGGAGAACCGGACCAUGAATT-3´ (DYNLT3_Oligo2) | Sigma-Aldrich | SASI_Hs02_00341758 | 50nM, 72h |
| Oligonucleotide | 5´- GAUUCAGAAUCCAACCGAATT-3´ (DYNLRB1) | Sigma-Aldrich | SASI_Hs01_00158744 | 50nM, 48h |
| Oligonucleotide | 5´-CUCAACUUUACCACCAAGUUATT-3´ (Spc24) | Sigma-Aldrich | (Xu et al., 2014b) | 50nM, 48h |
| Oligonucleotide | 5´-GGAUGAAGCAAGAGAUUUATT-3´ (NdeL1) | Sigma-Aldrich | SASI_Hs01_00228104 | 50nM, 48h |
| Oligonucleotide | 5´-GAGACAAGACUAUUAAGAUTT-3´ (LIS1) | Sigma-Aldrich | SASI_Hs01_00019092 | 50nM, 48h |
| Oligonucleotide | 5´-CAGCUAUUGCCACAGGGUUTT-3´ (HSET) | Sigma-Aldrich | SASI_Hs01_00185148 | 50nM, 48h |
| Oligonucleotide | 5´-GCUUGAAUCAGGCCAUCGATT-3´ (Nde1) | Sigma-Aldrich | SASI_Hs01_00074363 | 50nM, 48h |
| Oligonucleotide | 5´-GAUGCAUAUGAAGACUUUATT-3´ (DLIC2) | Sigma-Aldrich | SASI_Hs01_00014533 | 50nM, 72h |
| Oligonucleotide | 5´-CCAUUUCGCACGUAGCAGATT-3´ (HAUS6_Oligo1) | Sigma-Aldrich | SASI_Hs01_00105872 | 50nM, 48h |
| Oligonucleotide | 5´-CUAAUUGACUCUCUGGGUUTT-3´ (HAUS6_Oligo2) | Sigma-Aldrich | SASI_Hs01_00105873 | 50nM, 48h |
| Oligonucleotide | 5´-GUAUCUGAAUGCUUUGGUUTT-3´ (HAUS1_Oligo1) | Sigma-Aldrich | SASI_Hs01_00107510 | 50nM, 48h |
| Oligonucleotide | 5´-AAGGAUACCUCGCUAGCUAGUTT-3´ (HAUS1_Oligo2) | Sigma-Aldrich | (Einarson et al., 2004) | 50nM, 48h |
| Oligonucleotide | 5´-GAAACUCAGACUCCAGUUATT-3´ (DIC2) | Sigma-Aldrich | SASI_Hs01_00129736 | 50nM, 48h |
| Oligonucleotide | 5´- CAGACUUGGCCCAGUGUUUTT-3´ (survivin) | Sigma-Aldrich | SASI_Hs01_00052228 | 50nM, 24h |
| Oligonucleotide | 5´-GAUUUACGGGAAGAUUUGATT-3´ (CPAP) | Sigma-Aldrich | SASI_Hs01_00069692 | 50nM, 96h |
| Oligonucleotide | 5´-GAAUUGCAGCAGACUAUUATT-3´ (Hec1) | Sigma-Aldrich | SASI_Hs01_00138654 | 50nM, 24h |
| Oligonucleotide | 5´-CCUCUAAGGGAAUUCAGGATT-3´ (Borealin) | Sigma-Aldrich | SASI_Hs01_00153657 | 50nM, 48h |
| Oligonucleotide | 5´-CUACUACUUCUAACCUCCUTT-3´ (Scc1/Rad21) | Sigma-Aldrich | SASI_Hs01_00195799 | 50nM, 48h |
| Oligonucleotide | 5´-GCAAUACAGUGGGACAGUUTT-3´ (Myosin10) | Sigma-Aldrich | SASI_Hs01_00072460 | 50nM, 96h |
| Oligonucleotide | 5´-GCUGCACCAUGCCAAAUAATT-3´ (Sgo1) | Sigma-Aldrich | SASI_Hs01_00168960 | 50nM, 24h |

| | | | | |
|---|---|---|---|---|
| Oligonucleotide | 5´- GAGAAUGCCCAGUUAUUGATT-3´ (HAUS3) | Sigma-Aldrich | SASI_Hs01_00073692 | 50nM, 48h |
| Oligonucleotide | 5´-GAUUAAGGCUGUUAGUCUUTT-3´ (HAUS3) | Sigma-Aldrich | SASI_Hs01_00073693 | 50nM, 48h |
| Oligonucleotide | 5´-GAACUAGACUUGGUUAAUUTT-3´ (DHC) | Sigma-Aldrich | SASI_Hs01_00028998 | 50nM, 24h |
| Oligonucleotide | 5´- CAGUAGUGGCCAGGCUUCATT-3´ (CENP-T) | Sigma-Aldrich | (Chun et al., 2013) | 50nM, 48h |
| Oligonucleotide | 5´- GGAUGAAGUUAGAGAAAAUTT-3´ (MLL) | Sigma-Aldrich | (Ali et al., 2017) | 50nM, 24h |
| Oligonucleotide | 5´-CAGAUAAAGCGGAAGGCUCTT-3´ (CENP-W) | Sigma-Aldrich | (Chun et al., 2016) | 50nM, 72h |
| Oligonucleotide | 5´-AAGGUGAAUAUUGUGCCUGUCTT-3´ (Shp2_Oligo1) | Sigma-Aldrich | (Liu et al., 2012) | 50nM, 48h |
| Oligonucleotide | 5´-GGUUGCUACGGCUUAUCAUTT-3´ (Shp2_Oligo2) | Sigma-Aldrich | (Tsang et al., 2012) | 50nM, 72h |
| Oligonucleotide | 5´-GAAUCGUAUCUAUCUCACATT-3´ (ASURA/PHB2) | Sigma-Aldrich | (Takata et al., 2007) | 50nM, 24h |
| Oligonucleotide | 5´- UGGUUGAUGCAAGUGAAGATT-3´ (CENP-H) | Sigma-Aldrich | (Orthaus et al., 2006) | 50nM, 48h |
| Oligonucleotide | 5´- GUUGGCUAGAAUUGGGAAATT-3´ (Kif14_Oligo1) | Sigma-Aldrich | (Carleton et al., 2006) | 50nM, 48h |
| Oligonucleotide | 5´-GGCUCAGCAAGAGCUUUCUUCUCAATT-3´ (Kif14_Oligo2) | Sigma-Aldrich | (Xu et al., 2014a) | 50nM, 48h |
| Oligonucleotide | 5´- UUAGCAGUCACUCUUCCACTT-3´ (WDR5) | Sigma-Aldrich | (Ali et al., 2017) | 50nM, 48h |
| Oligonucleotide | 5´- CTAAGAGTTTGAAGTCTAATT-3´ (TAO1) | Sigma-Aldrich | (Draviam et al., 2007) | 50nM, 72h |
| Oligonucleotide | 5´-UGCUUUGUUGAACACAUCCTT-3´ (Nup88) | Sigma-Aldrich | (Bernad et al., 2004) | 50nM, 72h |
| Oligonucleotide | 5´- GAGAGAGGUCAAGCUGUGUGATT-3´ (ASB7) | Sigma-Aldrich | (Uematsu et al., 2016) | 50nM, 96h |
| Oligonucleotide | 5´- AAGATGGTCAAGAAGGCAGCATT-3´ (And-1_Oligo1) | Sigma-Aldrich | (Zhu et al., 2007) | 50nM, 72h |
| Oligonucleotide | 5´- GAUGGUCAAGAAGGCAGCATT-3´ (And-1_Oligo2) | Sigma-Aldrich | (Yoshizawa-Sugata and Masai, 2009) | 50nM, 72h |
| Oligonucleotide | 5´- ACGACUACAUUGAUAGUAAATT-3´ (Septin-7) | Sigma-Aldrich | (Zhu et al., 2008) | 50nM, 96h |
| Oligonucleotide | 5´-ACCUUGAUACUCAAUCAGGTT-3´ (ANKRD53) | Sigma-Aldrich | (Kim and Jang, 2016) | 50nM, 96h |
| Oligonucleotide | 5´-CGGACAAGCUGAACGAACATT-3´ (TRAMM) | Sigma-Aldrich | (Milev et al., 2015) | 50nM, 24h |
| Oligonucleotide | 5´-AAGGUGAAUAUUGUGCCUGUCTT-3´ (Septin-2_Oligo1) | Sigma-Aldrich | (Spiliotis et al., 2005) | 50nM, 48h |
| Oligonucleotide | 5´- GGUGAAUAUUGUGCCUGUCTT-3´ (Septin-2_Oligo2) | Sigma-Aldrich | (Kremer et al., 2005) | 50nM, 24h |
| Oligonucleotide | 5´-AAGACACAUAGUGGAUCUGUAUGTT-3´ (Seh1) | Sigma-Aldrich | (Zuccolo et al., 2007) | 50nM, 48h |
| Oligonucleotide | 5´-GGUCUGGCAUUACUACAGGAAGAAATT-3´ (CENP-Q_Oligo1) | Sigma-Aldrich | (Bancroft et al., 2015) | 50nM, 72h |
| Oligonucleotide | 5´- CAGAGUUAAUGACUGGGAAUAUUCATT-3´ (CENP-Q_Oligo2) | Sigma-Aldrich | (Bancroft et al., 2015) | 50nM, 72h |
| Oligonucleotide | 5´-AACUUCGGAAUUCUGCCUCUGTT-3´ (NF-1) | Sigma-Aldrich | (Park et al., 2013) | 50nM, 48h |
| Oligonucleotide | 5´-AAGAGGAAAGUGUAUUCGCAGTT-3´ (Nup107) | Sigma-Aldrich | (Zuccolo et al., 2007) | 50nM, 48h |
| Oligonucleotide | 5´-UAGUGAAUGUGGAAUGGAATT-3´ (Usp16) | Sigma-Aldrich | (Qian et al., 2016) | 50nM, 48h |
| Oligonucleotide | 5´- GAGCAGGTTGGCCACATTCTT-3´ (NDR1) | Sigma-Aldrich | (Oh et al., 2010) | 50nM, 96h |
| Oligonucleotide | 5´- GAUGUGCGGUUGUUCCUGGTT-3´ (GAK_Oligo1) | Sigma-Aldrich | (Shimizu et al., 2009) | 50nM, 48h |
| Oligonucleotide | 5´-AAGCUCAAGAUGUGGGGAGUG-3´ (GAK_Oligo2) | Sigma-Aldrich | (Lee et al., 2005) | 50nM, 72h |
| Oligonucleotide | 5´-GCGCUUAGAACGGAGUACUTT-3´ (HAUS7) | Sigma-Aldrich | SASI_Hs02_00350136 | 50nM, 48h |
| Oligonucleotide | 5´-GAAUUGACCUGAUCGUGUUTT-3´ (CENP-M) | Sigma-Aldrich | SASI_Hs01_00144699 | 50nM, 72h |
| Oligonucleotide | 5´-CUUUAUAAAUCAAUUGUUUTT-3´ (CENP-U) | Sigma-Aldrich | SASI_Hs01_00175574 | 50nM, 96h |
| Oligonucleotide | 5´-GACGUGUGCGGGAGAGUGATT-3´ (MST1) | Sigma-Aldrich | SASI_Hs01_00161455 | 50nM, 48h |

| | | | | |
|---|---|---|---|---|
| Oligonucleotide | 5´-GGUGUAAUGAUAUGUGCAUTT-3´ (PTEN) | Sigma-Aldrich | SASI_Hs01_00196478 | 50nM, 96h |
| Oligonucleotide | 5´-CUUUAGCAAAGAUGGAUAUTT-3´ (HAUS2) | Sigma-Aldrich | SASI_Hs01_00101146 | 50nM, 96h |
| Oligonucleotide | 5´-GACUGUUGCUGAGUUCAUCTT-3´ (CENP-N) | Sigma-Aldrich | SASI_Hs02_00322304 | 50nM, 72h |
| Oligonucleotide | 5´-CCGAGAAGGACGAGUUUGATT-3´ (Hsp72) | Sigma-Aldrich | SASI_Hs01_00051449 | 50nM, 72h |
| Oligonucleotide | 5´- GACAUGGAGAGGAAAGCCATT-3´ (HAUS5) | Sigma-Aldrich | SASI_Hs02_00347509 | 50nM, 48h |
| Oligonucleotide | 5´- GGAAGUUCAUCGUCUGAUUTT-3´ (HAUS4_Oligo1) | Sigma-Aldrich | SASI_Hs01_00022834 | 50nM, 48h |
| Oligonucleotide | 5´-GUGCUAUGAUCCUUAAGCUTT-3´ (HAUS4_Oligo2) | Sigma-Aldrich | SASI_Hs01_00022835 | 50nM, 48h |
| Oligonucleotide | 5´-CCAUGAAUUCAGUGAACUCTT-3´ (DYNLT1) | Sigma-Aldrich | SASI_Hs01_00096434 | 50nM, 48h |
| Oligonucleotide | 5´-GUCCUAUGCAGAUGACAAUTT-3´ (Rab5) | Sigma-Aldrich | SASI_Hs01_00097508 | 50nM, 48h |
| Oligonucleotide | 5´-AAGCAACUCGAAGAACAUCUCTT-3´ (CENP-I) | Sigma-Aldrich | (Liu et al., 2003) | 50nM, 72h |
| Oligonucleotide | 5´-GUCUAUCAGUGUCGAUUUATT-3´ (Dsn1) | Sigma-Aldrich | (Kim and Yu, 2015) | 50nM, 48h |
| Oligonucleotide | 5´-GAACCCUGGUAGGACUGCUUGGAAUTT-3´ (CENP-P) | Sigma-Aldrich | (McHedlishvili et al., 2012) | 50nM, 72h |
| Oligonucleotide | 5´-GGAGGACACUGCUAAGGGUTT-3´ (Zwint) | Sigma-Aldrich | (Zhang et al., 2015) | 50nM, 72h |
| Oligonucleotide | 5´-CAUGAGCUCUUUCUGUUUATT-3´ (Nsl1) | Sigma-Aldrich | (Kim and Yu, 2015) | 50nM, 48h |
| Oligonucleotide | 5´-GAAUCAUAAGGACUGUUCATT-3´ (Mis12) | Sigma-Aldrich | SASI_Hs01_00050622 | 50nM, 48h |
| Oligonucleotide | 5´-GCAUGUAUCUCUUAAGGAATT-3´ (KNL1) | Sigma-Aldrich | (Schleicher et al., 2017) | 50nM, 24h |
| Oligonucleotide | 5´-AAGGCAGACUCUACCAAAUGUTT-3´(Nup153) | Sigma-Aldrich | (Hahn et al., 2004) | 50nM, 48h |
